# Epidermal Growth Factor signaling acts directly and through a sedation neuron to depolarizes a sleep-active neuron following cellular stress

**DOI:** 10.1101/656512

**Authors:** Jan Konietzka, Maximilian Fritz, Silvan Spiri, Rebecca McWhirter, Andreas Leha, Sierra Palumbos, Wagner Steuer Costa, Alexandra Oranth, Alexander Gottschalk, David M. Miller, Alex Hajnal, Henrik Bringmann

## Abstract

Sleep is induced by sleep-active neurons that depolarize at sleep onset to inhibit wake circuits. Sleep-active neurons are under the control of homeostatic and allostatic mechanisms that determine sleep need. However, little is known about the molecular and circuit mechanisms that translate sleep need into the depolarization of sleep-active neurons. During many conditions in *C. elegans* sleep induction requires a sleep-active neuron called RIS. Here, we defined the transcriptome of RIS to discover that genes of the Epidermal Growth Factor Receptor (EGFR) signaling pathway are expressed in RIS. With cellular stress, EGFR activates RIS, and RIS induces sleep. Activation of EGFR signaling in the ALA neuron has previously been suggested to promote sleep independently of RIS. Unexpectedly, we found that ALA activation promotes RIS depolarization. Our results suggest that ALA is a sedating neuron with two separable functions. (1) It inhibits specific wakefulness behaviors independently of RIS, (2) and it activates RIS to induce sleep. Whereas ALA plays a strong role in surviving cellular stress, surprisingly, RIS does not. In summary, EGFR signaling can induce sleep-active neuron depolarization by an indirect mechanism through activation of the sedating ALA neuron that acts upstream of the sleep-active RIS neuron as well as through a direct mechanism using EGFR signaling in RIS. Sedation rather than sleep appears to be important for increasing survival following cellular stress, suggesting that sedation and sleep play different roles in restoring health.

**Highlights:**

- The transcriptome of the sleep-active RIS neuron reveals the presence of the EGFR signaling machinery
- EGFR activates RIS directly upon cellular stress to induce sleep bouts
- In parallel, EGFR activates RIS indirectly through the sedating ALA neuron
- Sedation rather than sleep bouts support survival following cellular stress

## Introduction

Sleep is a vital state in animals, and thus mechanisms exist that tightly monitor and fulfill sleep need[1]. Circadian, homeostatic, and allostatic mechanisms control when and how much an animal sleeps[2, 3]. The circadian rhythm controls the timing of sleep. In the absence of a circadian rhythm, the majority of sleep will still take place, but sleep will not be confined to a particular time of the day[2, 4, 5]. Sleep homeostasis translates sleep need caused by prior wakefulness into increased sleep drive. Homeostasis can act acutely to ensure that sleep is maintained by detecting arousal during sleep and promoting sleep maintenance or a return to sleep[6, 7]. Also, sleep homeostasis can integrate over prolonged wake activity to match the time spent awake to subsequent sleep. Sleep homeostasis involves somnogens, substances that accumulate as a function of wakefulness and that promote sleep. An example for a somnogen is adenosine, which is released from astrocytes as a function of the activity of neighboring neurons[8–10]. Also, glutamate receptor activity increases in a subset of sleep-promoting neurons during wakefulness, thus leading to an increase in their excitability and an increase in sleep pressure. Sleep in turn reduces the plasticity of these neurons thus releasing sleep pressure[11]. Allostatic mechanisms increase sleep in response to an insult, thus supporting the restoration of physiological homeostasis. In humans, excessive daytime sleepiness is a pathological condition that is characterized by a general lack of energy during the day despite normal or even increased nighttime sleep. This reduced behavioral activity during wakefulness and its associated increase in sleep is often described as sedation, drowsiness, lethargy, somnolence, or excessive daytime sleepiness, but is ill defined at the mechanistic level [12–14].

Circadian, homeostatic, and allostatic mechanisms drive sleep by controlling the depolarization of sleep-active sleep-promoting neurons. These neurons depolarize at sleep onset to release inhibitory neurotransmitters, peptides and GABA, onto wakefulness and arousal circuits to dampen their activity. Inhibition of wake circuits leads to the cessation of voluntary movements and the dampening of afferent sensory stimulus processing. Sleep reversibility, i.e. waking up, involves activation of arousal circuits and inhibition of sleep-active neurons, whereas sleep homeostasis involves an increased activation of sleep-active neurons during rebound sleep following sleep deprivation[15, 16].

Sleep-active neurons were first identified in mammals, where several populations of these neurons exist. Well characterized are sleep-active neurons in the preoptic area of the hypothalamus. These neurons inhibit the ascending arousal system through GABA and neuropeptides, including galanin. Sleep deprivation leads to increased depolarization of sleep-active neurons, indicating that they transduce increased sleep pressure derived from upstream driver circuits[16–19]. Additional populations of sleep-active neurons are found in the basal forebrain, lateral hypothalamus, and the parafacial zone of the brain stem. It is likely that these various populations act in concert to induce sleep[17, 20]. Multiple signaling molecules have been proposed to promote sleep during various conditions, including during allostatic sleep pressure. Epidermal Growth Factor (EGF) signaling is involved in growth and healing processes as well as in sleep control. Infusion of the EGF receptor (EGFR) ligand tumor necrosis factor alpha (TNF-α), which is rhythmically expressed in the suprachiasmatic nucleus, or of EGF into the cerebrospinal fluid reduces behavioral activity and increases sleep[21–23]. Upon infection, proinflammatory cytokines, including interleukin-1 beta (IL1) and TNF-α, are released that affect neuronal excitability, synapse composition and induce sedation and increased sleep[9, 24]. TGF-α infusion broadly reduces behavioral activity including grooming, feeding, exploring, and locomotion, indicating a general inhibition of active behaviors that has been described to be similar to “sickness behavior”, in which animals become lethargic and anorexic[23]. Sedation and subsequently increased sleep in mammals after infection correlates with increased recuperation and decreased mortality. For example, sleep is a positive predictor for recuperation from influenza infection in rabbits[9, 24, 25].

As in mammals, sleep-active neurons are also found in invertebrates. In *Drosophila*, several populations of neurons were found to promote sleep, including neurons in the mushroom body and dorsal paired medial neurons[26, 27]. The dorsal fan-shaped body contains neurons that are similar to mammalian sleep-active neurons in that they can actively induce sleep and their ablation reduces sleep. These neurons increase their activity after sleep deprivation suggesting that they act downstream of sleep homeostasis[11, 28, 29]. Overexpression of TGF-α in *Drosophila* leads to an increase in sleep, whereas inhibition of EGFR caused a shortening of sleep bouts. TGF-α is released from neurons of the pars intercerebralis, which is analogous to the hypothalamus in vertebrates, to activate EGFR signaling in several neurons, including those of the tritocerebrum, leading to the activation of extracellular signal-regulated kinase (ERK). EGFR signaling likely leads to alterations of neuronal activity, but how EGF signaling leads to the induction of sleep is not known[30].

*C. elegans* possess a sleep-active and sleep-inducing neuron called RIS, which is crucially required for sleep during many physiological conditions. Like its mammalian counterparts, this neuron activates at the onset of sleep bouts and actively induces sleep[31–34]. RIS increases its activity after sleep deprivation to induce rebound sleep[33, 35, 36]. Allostatic load caused by cellular stress leads to EGF release, which activates a second neuron, called ALA, through the EGFR and downstream phospholipase C (PLC) pathway. ALA induces increased sleep by releasing multiple neuropeptides that alter neuronal physiology [37–39]. ALA has been proposed to act in parallel to the RIS sleep circuit to increase sleep following cellular stress. It has been suggested that ALA can inhibit different behaviors, but the mechanism by which ALA induces dynamic sleep bouts following EGF release is not understood[39, 40]. Mutant worms that have reduced sleep after heat shock due to the loss of ALA function have impaired survival, indicating that allostatic sleep-enhancing mechanisms are protective not only in mammals but also in invertebrates [41, 42]. Thus, across species, allostatic sleep pressure leads to the release of somnogens that increase protective sleep. In mammals, flies, and worms, EGF release signals sleep need, suggesting that EGFR signaling presents an evolutionarily ancient pathway for sleep induction. However, the mechanisms by which release of somnogens such as EGF leads to the depolarization of sleep-active neurons is not well understood in any system. Comprehending allostatic sleep need requires understanding two interconnected behaviors, the reduced behavioral activity during wakefulness that is typical of sedation, as well as increased sleep caused by the increased depolarization of sleep-active neurons.

Here we aimed to understand the control of the sleep-active RIS neuron by upstream mechanisms through analyzing its molecular content. We obtained the transcriptome of RIS, genetically screened RIS-expressed genes for sleep phenotypes, and analyzed the mechanisms by which these genes control RIS depolarization. This analysis identified a mechanism explaining how EGFR signaling controls sedation and sleep upon cellular stress by directly activating both, the sedating ALA neuron as well as the sleep-bout inducing RIS neuron.

## Results

### Transcriptome analysis reveals the presence of the EGFR signaling machinery in RIS

To find molecular pathways controlling RIS, we identified the transcriptome of this neuron. To obtain high-quality transcriptome data of RIS, we generated RIS transcriptomes by two complementary methods. The first approach was to extract the RIS transcriptome from single-cell data sets and the second approach was to obtain the RIS transcriptome by RNA-seq of FACS-isolated cells. To identify the RIS transcriptome from single-cell data, we used available transcriptomes of all cells of L2 larvae of *C. elegans* that have been profiled using single-cell combinatorial indexing RNA sequencing (sci-RNA-seq)[43]. To identify the transcriptome cluster corresponding to RIS within the neuronal sci-RNA-seq clusters, we used our previous observations that only RIS strongly and specifically expresses *flp-11* neuropeptides [32]. One cluster stood out among the transcriptomes in that it showed high counts for *flp-11* expression for every cell that was assigned to this cluster (Figure 1). Differential expression analysis versus all other neurons and all other cells identified 66 and 381 significantly changed genes, respectively (Table S1-S2). The most strongly enriched gene in RIS was *flp-11*, with an enrichment of 157-fold and 588-fold, respectively. To generate an RIS transcriptome from isolated cells we FACS-sorted dissociated cells derived from L2 larvae expressing the red-fluorescent mKate2 protein specifically in RIS using the *flp-11* promoter [44, 45]. We obtained an RIS transcriptome containing 4371 genes expression of which was significantly changed (Table S3). Again, *flp-11* was among the most highly enriched of all genes (enrichment of 891-fold).

**Figure 1.**
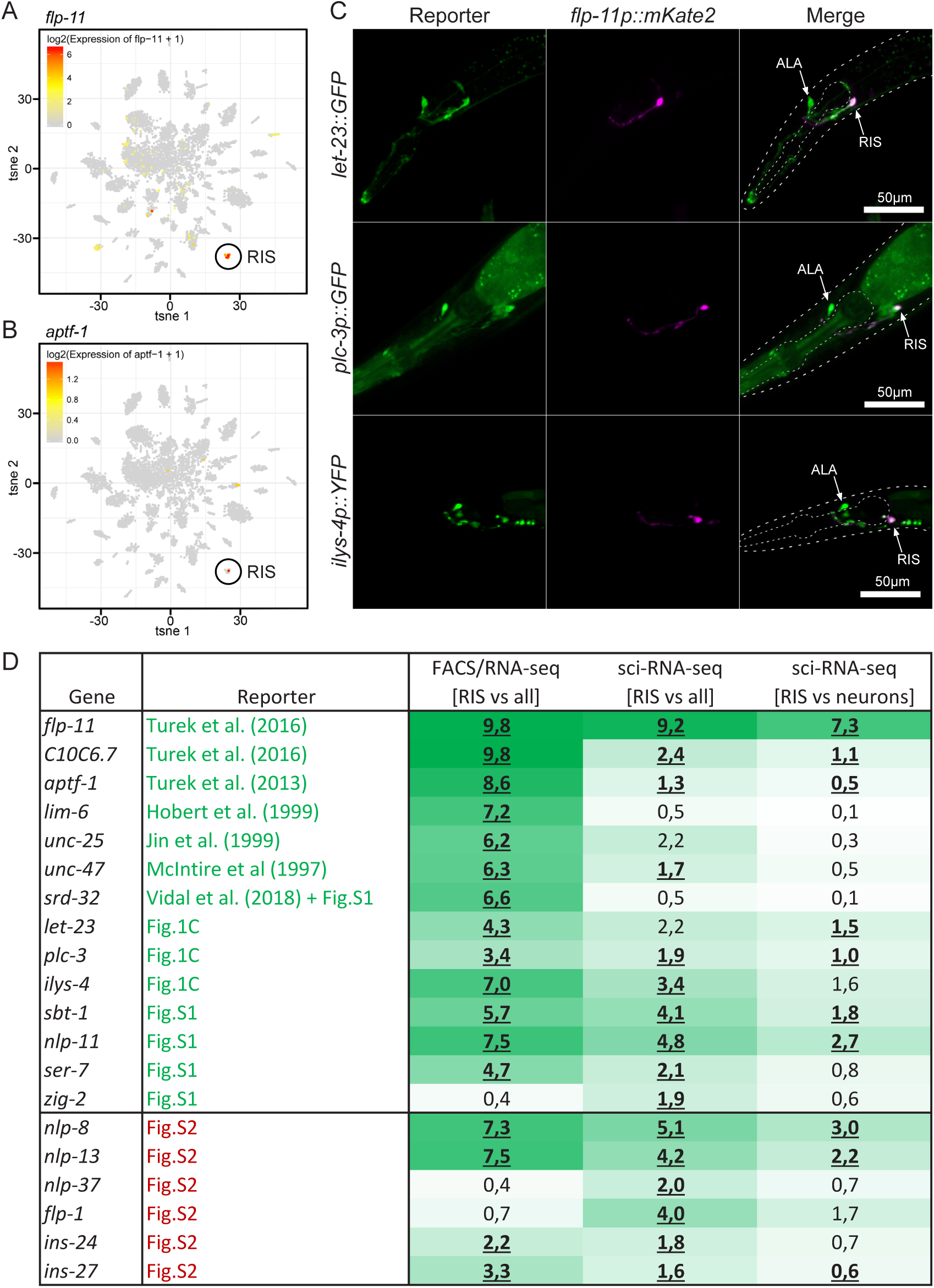
Transcriptome analysis of the sleep-active RIS neuron. (A-B) Identification of RIS from sci-RNA-seq data. tsne-plots of all neuronal cells were color coded for log2 expression values of (A) *flp-11* or (B) *aptf-1*. (C-D) Validation of RIS enriched genes using fluorescent transgene transcriptional or translational reporters. (C) Example micrographs for *let-23::GFP*, *plc-3p::GFP*, *ilys-4p::GFP*, and their co-localization with *flp-11p::mKate2*. Dashed lines display the outlines of the head and pharynx (anterior is left, dorsal is up). ALA and RIS are indicated with white arrows. Scale bar is 50µm. (D) Table summarizing genes tested for fluorescence-reporter expression in RIS via colocalization of *flp-11p::mKate2* and comparison of RIS transcriptomes obtained by either bulk sequencing of FACS-isolated cells or sci-RNA-seq. Enrichment is displayed as log2FC and color coded with darker green color indicating more enrichment in RIS. Significantly enriched genes are displayed as bold and underlined. For statistical comparison a likelihood ratio test was used, adjusted for multiple testing using Benjamini-Hochberg (α = 5% for FACS/RNA-seq, α = 10% for sci-RNA-seq).

To validate the RIS transcriptomes, we used 20 available reporter transgenes expressing fluorescent proteins. For 14 of these genes we could see reporter gene expression in RIS, 7 of these genes were reported previously to localize to RIS (the GABA biosynthetic gene *unc-25*[46], the vesicular GABA transporter gene *unc-47*[47], the transcription factor genes *lim-6* [48] and *aptf-1* [31], *flp-11* and *C10C6.7* [32], and *srd-32* [49]). Reporter lines showed RIS expression in an additional seven genes that were not previously reported to be expressed in RIS (*let-23*, *plc-3, ilys-4*, *sbt-1, nlp-11, ser-7, zig-2*) (Figure 1C, Figure S1). Six transgenes did not show expression and could be either false positives in the transcriptome or false negatives due to the limitations of reporter transgenes (Figure S2). The transcriptome derived from FACS sorting contained most of the genes expected to be expressed in RIS based on reporter gene analysis, but appeared to contain also more broadly expressed genes. The single-cell RIS transcriptome contained only the most specifically expressed genes. In summary, we have identified the RIS transcriptome and provide lists of differentially expressed genes with different levels of stringency. This transcriptome resource thus provides an entry point into studying the molecular biology of RIS.

To identify sleep-controlling genes that regulate RIS, we performed a behavioral screen based on the RIS transcriptome. We focused on 106 genes that were strongly enriched and for which mutations were available. We quantified sleep of these mutants by filming their behavior inside agarose hydrogel microcompartments. We used starved arrested L1 larvae for this screen because *C. elegans* displays robust RIS-dependent sleeping behavior under these conditions[33]. The fraction of time spent asleep was extracted using automatic image analysis based on frame subtraction[50]. As many of the mutations were obtained from mutagenesis screens, we backcrossed alleles that produced a significant phenotype in the primary screen and tested them again. We obtained seven genes whose mutation changed sleep amount significantly and more than 50% from wild type (Figure 2A). The four genes that reduced sleep had already been described to control sleep (*aptf-1*, *flp-11*, *goa-1*, *frpr-3*)[31, 32, 51]. We obtained three genes whose mutation caused increased sleep. *nhr-128* and *ilys-4* showed almost a doubling of time spent in sleep. A dominant gain-of-function of *let-23*, (*let-23(sa62), let-23(gf)*[52]), showed almost a tripling of mean sleep time (Figure 2A).

**Figure 2.**
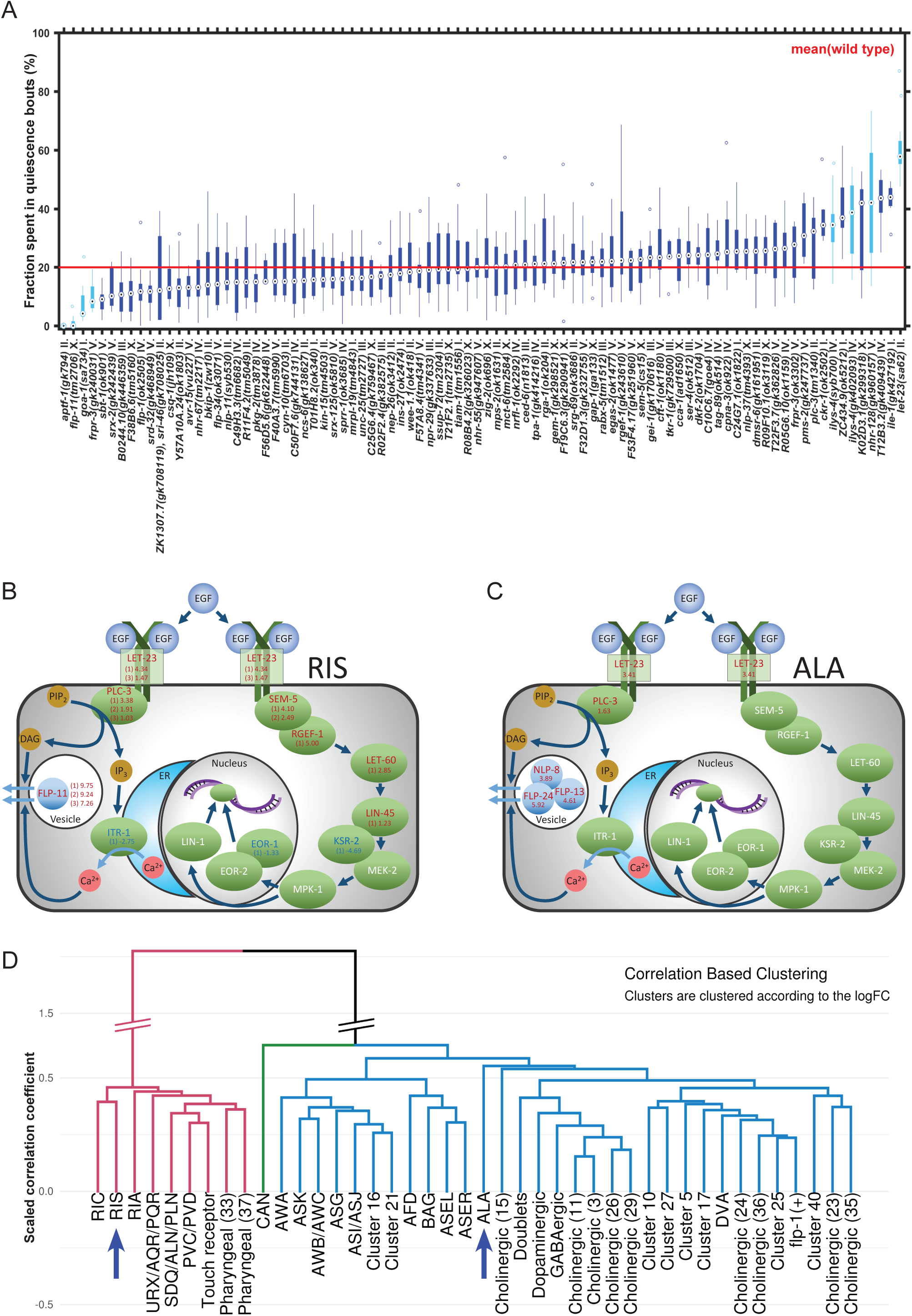
The EGFR signaling machinery is expressed in both ALA and RIS neurons. (A) L1 arrest sleep screen for mutants of genes that are enriched in RIS. Every allele screened is represented by a blue boxplot (alleles are on the x-axis, fraction (%) spent in locomotion quiescence on the y-axis), wild-type mean is displayed as a red line. Alleles that deviated more than 50% of the wild type mean and were significantly different to the respective control data are indicate in turquoise, p < 0.05, Wilcoxon signed-rank test. The strongest increase of sleep is seen in a gain-of-function mutation of *let-23*. (B-C) Both ALA and RIS express EGFR pathway components. Shown are enrichments of canonical EGFR signaling components in RIS (B) and ALA (C). Proteins (green) or neuropeptides (light blue), which were found to be significantly differentially expressed in at least one of the transcriptomes are indicated (enriched is red, de-enriched is blue, no significant change is white). (B) Gene expression changes are displayed as log2FC ((1) indicates RIS vs. all, FACS/RNA-seq; (2) indicates RIS vs. all, sci-RNA-seq; (3) indicates RIS vs. neuron, sci-RNA-seq). (C) Gene expression changes are displayed as log2FC (ALA vs. neuron, sci-RNA-seq). PIP2: phosphatidylinositol 4,5-bisphosphate, IP3: inositol trisphosphate, DAG: diacylglycerol, ER: endoplasmic reticulum. Likelihood ratio test, adjusted for multiple testing using Benjamini-Hochberg, α = 5% for FACS/RNA-seq, α = 10% for sci-RNA-seq. (D) Despite an overlap of expression of EGFR signaling components, RIS and ALA are divergent in overall gene expression. Correlation-based clustering of all neuronal clusters identified from single-cell sequencing. Blue arrows indicate the clusters corresponding to RIS and ALA. Correlation coefficient was rescaled to display strong correlation as small distances (0 corresponds to perfect correlation (correlation coefficient = 1), 2 corresponds to perfect negative correlation (correlation coefficient = −1), 1 corresponds to uncorrelated (correlation coefficient = 0).

*let-23* encodes the sole EGF receptor in *C. elegans*. EGF signaling activates intracellular pathways, including PLC controlling cellular excitability, *vav-1* Rho GEF, and Ras controlling gene expression[53]. We hence looked at whether typical EGFR signaling components are also expressed in RIS. Canonical EGF signaling genes were expressed in RIS and several components of the PLC and Ras pathway were enriched (Figure 2B, *vav-1* an *rho-1* were not significantly enriched) [53]. The presence of the EGF signaling machinery also in RIS suggests that not only ALA[37], but also RIS is involved in the EGF response to cellular stress. Hence, we compared RIS and ALA transcriptomes. For this comparison, we extracted the ALA transcriptome from the single cell data set, this time based on the expression of *flp-24*, *flp-13*, and *flp-7* [38, 39]. ALA transcriptomes could be clearly identified (methods). We calculated a list of differentially expressed genes for ALA by comparison to all other neurons (Table S4). ALA also expressed canonical components of the EGF intracellular signaling pathway but only PLC3 was enriched (Figure 2C)[37, 39]. Pairwise correlations of differential expression for all discernable neuronal transcriptomes and hierarchical clustering revealed a relatively low degree of similarity in gene expression for RIS and ALA (Figure 2D). Thus, ALA and RIS both express EGFR signaling components but otherwise their overall molecular contents are different suggesting that both neurons are activated by EGF but are functionally divergent.

### EGFR signaling acts in both ALA and RIS to induce sleep after cellular stress

*let-23* promotes sleep and is expressed in two neurons, ALA and RIS, that have both been implicated in sleep induction, with RIS acting in physiological sleep and ALA acting in stress-induced sleep[31, 33, 37, 40, 41]. To test the contributions of these two neurons to EGF-induced sleep, we ablated these neurons genetically in the *let-23gf* mutant and quantified starvation-induced sleep during L1 arrest. To functionally ablate ALA, we used a deletion in the homeodomain transcription factor gene *ceh-17*, which is required for expression of EGF signaling in ALA[37]. For RIS ablation we used a strain that expresses the apoptosis inducer *egl-1* specifically in RIS[33]. Ablation of ALA suppressed the increased sleep phenotype partially, and ablation of RIS removed all sleep behavior (Figure 3A). Thus, *let-23gf*-induced sleep during larval arrest involves both ALA and RIS. ALA accounts for much but not all of the increased sleep, suggesting that ALA plays a major role in increasing sleep but is not the sole factor increasing sleep upon EGFR activation. Virtually all sleep bouts required RIS, suggesting that RIS is the actual inducer of EGFR-triggered sleep bouts during L1 arrest. These results are consistent with a model in which EGF receptor activation promotes sleep through two pathways, the first could activate RIS directly and the second could activate RIS indirectly through ALA.

**Figure 3.**
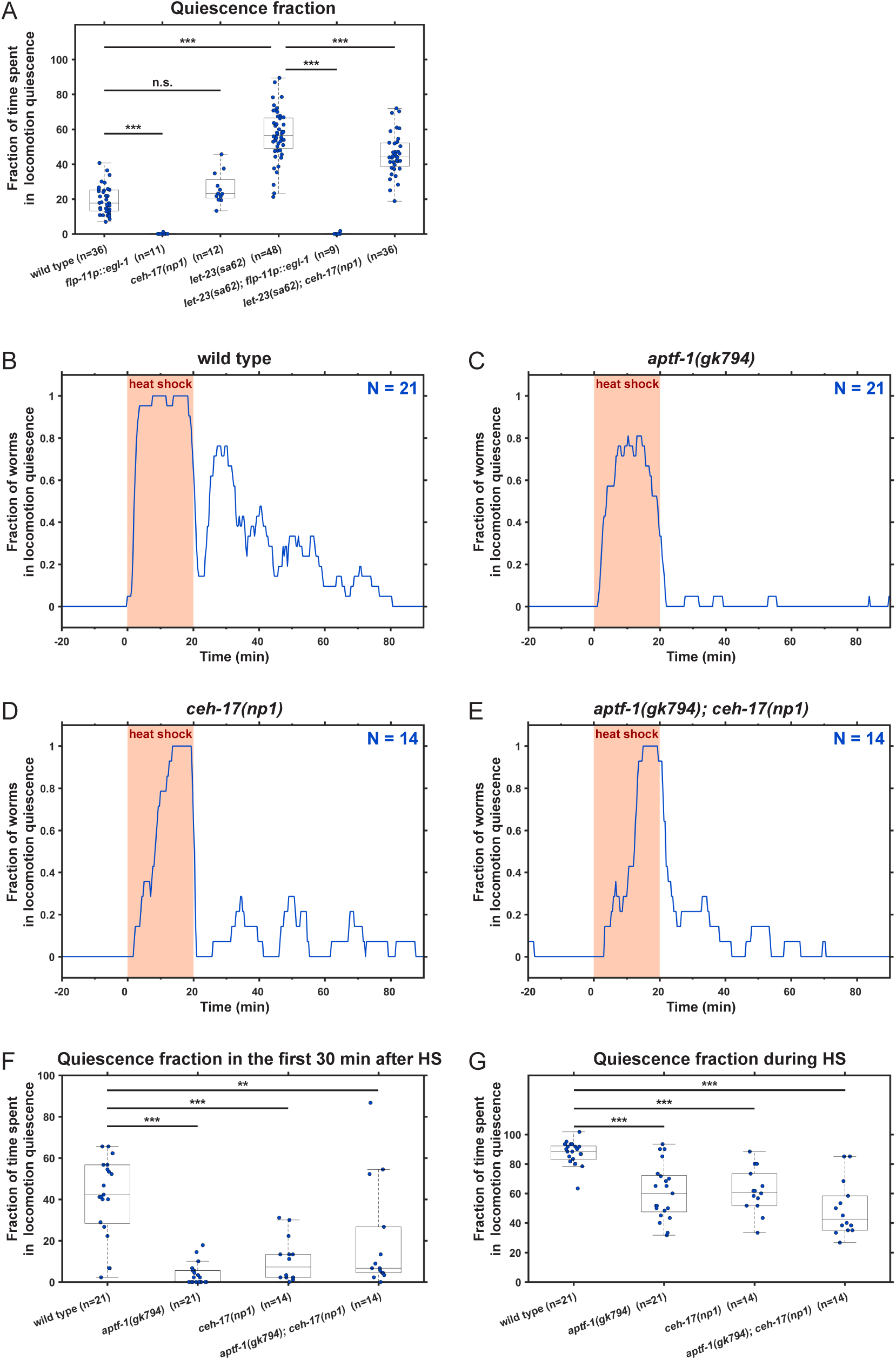
ALA and RIS are required for sleep after EGFR activation and cellular stress. (A) ALA and RIS are required for sleep caused by *let-23gf* during L1 arrest. Fraction of time spent in sleep during L1 arrest was measured by quantifying locomotion quiescence. *ceh-17lf* partially suppressed the increase of sleep caused by *let-23gf*. RIS-ablation suppressed virtually all sleeping behavior. (B-E) ALA and RIS are required for sleep induced by cellular stress. The fraction of worms is shown vs. time, with the time during which the heat shock (37°C) was applied is indicated in orange. N is number of worms; three biological replicates were performed for each genotype. (B) Wild-type worms immobilized during heat shock and a series of consecutive quiescence bouts followed during the 60 minutes after the heat shock. (C) *aptf-1lf* showed reduced movement quiescence during the heat shock and almost no quiescence was seen following the heat shock. *ceh-17lf* mutants took longer to immobilize during the heat shock and sleep bouts were reduced following the heat shock. (E) *aptf-1lf*; *ceh-17lf* double mutation reduced quiescence, albeit not as strongly as *aptf-1lf* alone. (F-G) Quantification of locomotion quiescence. *** denotes statistical significance with p < 0.001, ** denotes statistical significance with p < 0.01, Wilcoxon signed-rank test.

The expression of the EGFR in ALA and RIS suggested that, upon cellular stress, EGF activates not only ALA as shown previously[37, 41], but also RIS to induce sleep. To test this idea, we induced cellular stress and measured locomotion behavior afterwards in wild-type and in mutants in which RIS and ALA were genetically ablated (*aptf-1(-)* and *ceh-17(-)*, respectively). To induce cellular stress, we placed adult worms in microfluidic compartments and delivered a heat shock for 20 minutes at 37°C using a Peltier element (Figure S3). This setup allowed the continuous quantification of sleep behavior before, during, and after the heat shock. Wild-type worms showed no detectable sleep behavior before the heat shock. During the heat shock, worms acutely immobilized. After the heat shock, worms showed a succession of sleep bouts that appeared highly rhythmic and were most pronounced during the first 20 minutes and then slowly decayed over one hour. RIS ablation reduced quiescence during the heat shock by about 30%, but virtually abolished locomotion quiescence after heat shock. ALA ablation slowed down the immobilization during the heat shock and led to a reduction of sleep following the heat shock by about 80% similar to previous reports[37, 41, 42]. Ablation of both RIS and ALA led to an extension of the immobility caused by the heat shock, suggesting that double ablation perhaps caused some unspecific quiescence, but locomotion quiescence levels eventually decreased to a similar level as observed after *aptf-1(-)*(Figure 3B-G). Thus, in addition to the ALA neuron that has been long studied in stress-induced sleep [37, 41, 42], the RIS neuron is crucially required for sleep bout induction following cellular stress. This requirement of RIS to induce sleep bouts following cellular stress is consistent with a recent report that used chemogenetic inhibition of RIS[54].

The neuronal ablation experiments showed that both ALA and RIS are required to promote sleep following EGFR activation. EGFR signaling has been shown previously to act in ALA[37]. To test whether the EGFR is also required in RIS following cellular stress, we knocked out *let-23* specifically in RIS and quantified sleep following the heat shock. We used a conditional allele of *let-23*, in which three exons containing the critical kinase domain are flanked by *frt* sites and can thus be removed through FLPase-induced recombination[55–57]. We expressed FLPase in RIS using a promoter (*unc-47*) that does not express in ALA, which reduced the expression of *let-23* specifically in RIS based on GFP expression. Specific deletion of *let-23* in RIS led to a reduction of sleep by more than half compared with the parental strains (Figure 4). Thus, EGF signaling is required in both ALA and RIS to promote sleep following heat shock. These results suggest a model in which EGF is released following cellular stress and activates both ALA and RIS, which act concertedly to induce sleep. The consistently stronger effect of RIS impairment on sleep bouts compared with ALA impairment suggests that ALA might act upstream of RIS.

**Figure 4.**
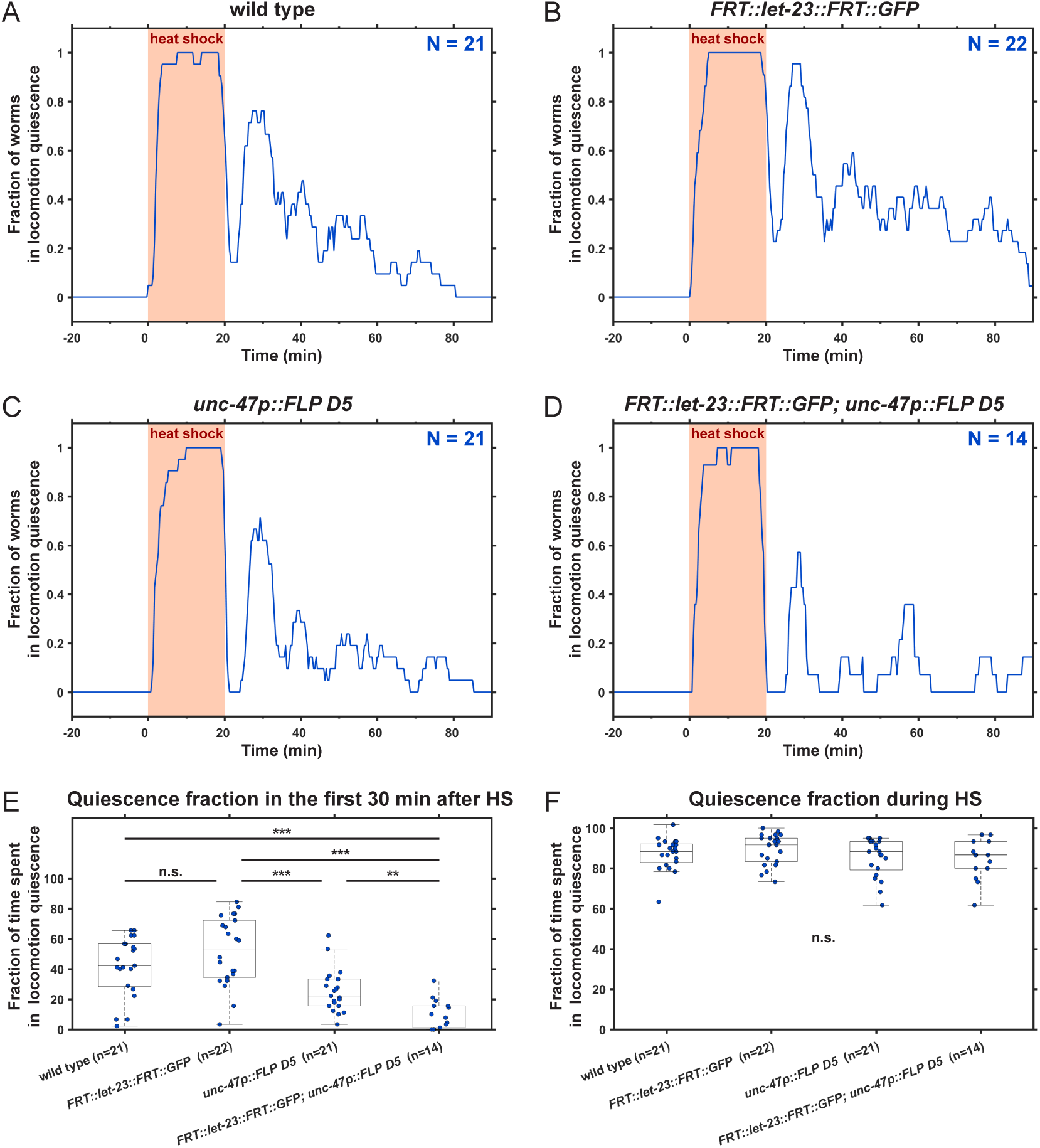
EGFR is required in RIS to increase sleep after cellular stress. (A-D) RIS-specific knockdown of *let-23* reveals a role for EGFR in RIS following cellular stress, in addition to the known role of EGFR in ALA[37]. The heat shock (37°C) is indicated in orange. N indicates number of worms; three biological replicates were performed for each genotype. (A) is the same data as in Figure 3B. (B-C) behavior of the parental strains following a heat shock. (B) The conditional allele of *let-23*, *FRT::let-23::FRT::GFP*, was created using CRISPR/Cas9. (C) The recombinase (FLPase) was expressed in GABAergic neurons, *unc-47p::FLP D5*, which include RIS, but no expression was detectable in ALA with this promoter (data not shown). (D) Worms with a conditional knockdown of *let-23* in RIS, *unc-47p::FLP D5; FRT::let-23::FRT::GFP,* displayed reduced quiescence. (E-F) Quantification of locomotion quiescence during and after the heat shock in the conditional strain and in the parental controls. *** denotes statistical significance with p < 0.001, Wilcoxon signed-rank test.

### Cellular stress and EGF signaling depolarize RIS and ALA

Calcium imaging and optogenetic manipulation of RIS and ALA have suggested that these neurons act through depolarization [31, 38]. To test whether cellular stress and EGF depolarize RIS and ALA, we measured calcium activity in these neurons after heat shock or overexpression of the EGF gene *lin-3*. We first looked at RIS and applied a heat shock inside the microfluidic device and measured calcium activity using GCaMP. Upon temperature increase, RIS activated strongly while the animal immobilized, which is consistent with a previously identified increase of RIS during temperature increase [58]. After the heat shock, calcium transients were strongly correlated with sleep bouts. Calcium transients occurred rhythmically following the heat shock. Typically, three to four consecutive RIS transients and sleep bouts lasting each for about 12 minutes were observed, with the first transient being the strongest and subsequent transients displaying reduced intensity until the succession of transients ceased after about one hour. Correlation analysis showed that RIS activated specifically during sleep bouts (Figure 5A). ALA ablation resulted in a strong reduction of RIS activity (Figure 5B).

**Figure 5.**
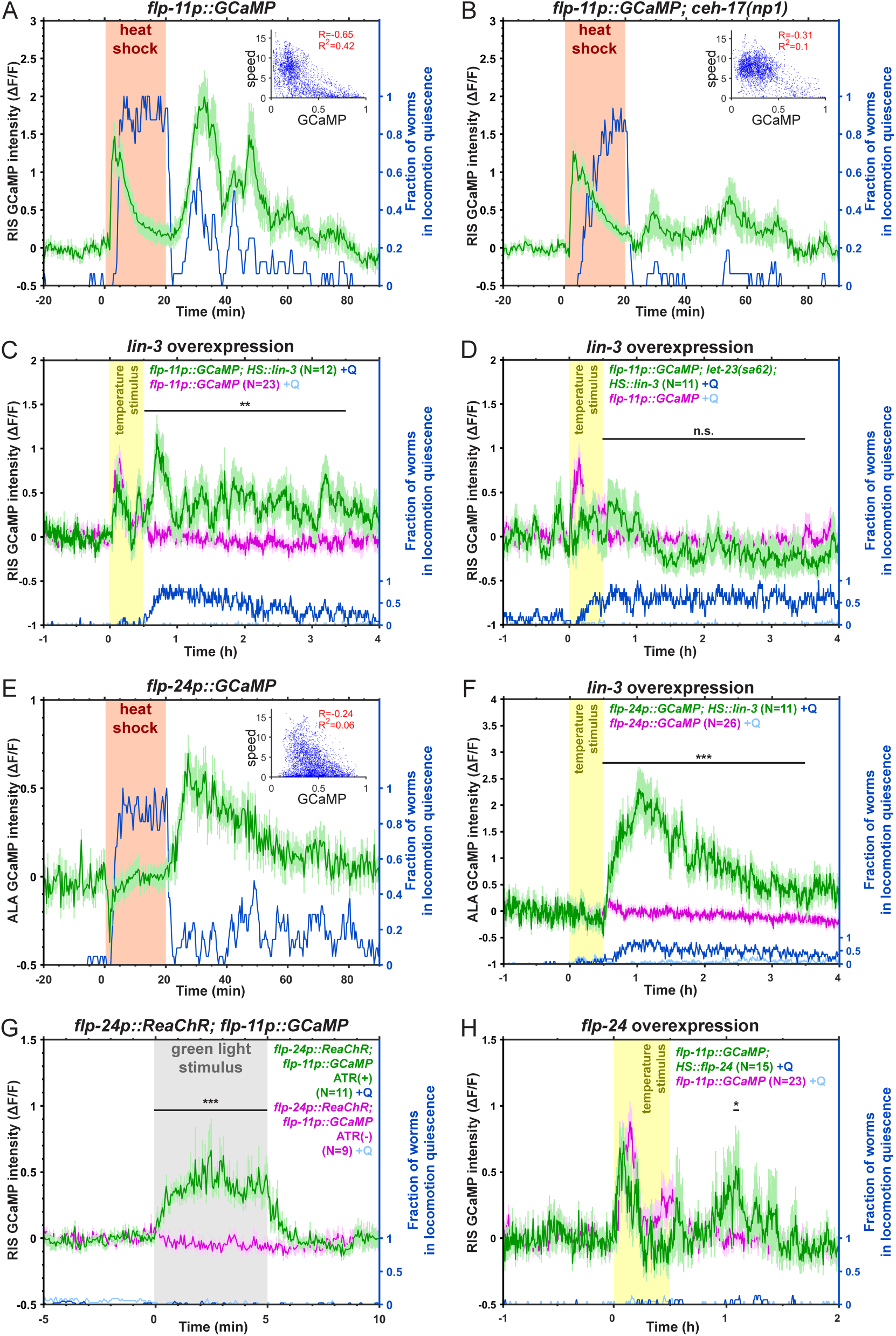
Heat shock and EGF activate RIS and ALA, and ALA acts upstream of RIS. (A) Calcium activity of RIS following a heat shock (37°C, orange). *flp-11p::GCaMP* intensities are shown in green, and the fraction of worms in locomotion quiescence is shown in blue. The insert shows a correlation of normalized, smoothed GCaMP intensities with speed (µm/s) for the first 60 min after the heat shock, with Spearman’s rank correlation coefficient. Calcium activity peaks during the first part of the heat shock and shows several transients following the heat shock. RIS calcium transients correlate with locomotion quiescence. (B) *ceh-17lf* reduces RIS calcium activity following the heat shock. (C) EGF over expression induces RIS calcium activity. Overexpression is induced by a temperature increase (30°C, yellow). Control *flp-11p::GCaMP* (without EGF overexpression transgene) intensity (magenta) ±SEM, according fraction of worms in movement quiescence (light blue). (D) EGF over expression in *let-23gf* does not further increase RIS calcium activity. *let-23gf* leads to movement quiescence already before the heat shock, while no increase in GCaMP activity can be seen. (E) A heat shock causes subsequent ALA calcium activation. GCaMP activity does not increase during the heat shock but after the heat shock. Neural activity and locomotion quiescence do not correlate well. (F) EGF overexpression by a heat-shock promoter and temperature increase induces massive ALA activation. (G) Optogenetic activation of ALA by green light (indicated in grey) causes RIS calcium activation (green). Control (without retinal) *flp-11p::GCaMP* intensity (magenta). Movement quiescence is shown in light blue. (H) Overexpression of *flp-24* by a heat-shock promoter and temperature increase (30°C, yellow) induces RIS calcium transients. Error is ± SEM, *** denotes statistical significance with p < 0.001, ** denotes statistical significance with p < 0.01, * denotes statistical significance with p < 0.05, Wilcoxon signed-rank test.

To test for the effects of EGF upon RIS activation, we overexpressed this signaling protein using a heat-shock promoter. We induced expression with a temperature increase that is not sufficient to trigger subsequent sleep. Overexpression of EGF induced immobility as reported previously [37] and led to a strong increase of RIS calcium activity (Figure 5C). RIS calcium imaging in *let-23(gf)* mutant animals showed that RIS is already active during baseline condition before the heat shock and cannot be activated much further following EGF overexpression (Figure 5D). We next measured calcium activity of ALA. This neuron showed an increase in calcium activity following the heat shock that correlated with the time during which sleep bouts occurred. However, ALA activity did not correlate well with the sleep state of the animal, i.e. correlation analysis showed that ALA did not activate specifically during sleep bouts but are more broadly associated with the time during which sleep bouts occur (Figure 5E). Overexpression of EGF increased the calcium activity of ALA substantially, and the calcium increase was much stronger than that following a heat shock (Figure 5F).

These results show that cellular stress and EGF increase calcium activity of RIS and ALA. Intriguingly, the activation kinetics of these neurons differed. ALA activity correlated with the time during which sleep bouts occurred, but calcium activity did not correlate strongly with the actual sleep state. By contrast, RIS activation transients directly correlated with the occurrence of sleep bouts. The different calcium kinetics of ALA and RIS suggest that these neurons act by different mechanisms, with ALA inducing sleep bouts indirectly and RIS inducing sleep bouts directly. Together with the reduction of RIS calcium transients in the absence of ALA these kinetic changes suggest that ALA activates RIS to induce sleep bouts.

It has been shown that ALA can act independently of RIS to induce quiescence [40]. Our results suggest that ALA can also act upstream of RIS to induce dynamic sleep bouts. ALA has been proposed to induce sleep by calcium-induced secretion of multiple neuropeptides that may act by a diffusional mechanism, but ALA also has been shown to control locomotion behavior and sleep through synaptic mechanisms [38, 39, 59, 60]. To test whether ALA activates RIS, we optogenetically activated ALA and measured the response of RIS using calcium imaging. We expressed the red light-activated channelrhodopsin variant ReaChR specifically in the ALA neuron and GCaMP in the RIS neuron. ALA was activated using green light for 5 minutes and RIS activity was followed using fluorescence microscopy. RIS activated upon optogenetic ALA activation and its activity decreased back to baseline levels once the ALA stimulation ceased (Figure 5G).

Because ALA has been shown to act through neuropeptides to induce quiescence, we next overexpressed *flp-24*, a key ALA-expressed sleep-promoting neuropeptide [39]. *flp-24* was expressed from a heat shock-inducible promoter and a mild temperature increase that does not induce sleep by itself as above. Consistent with a role of this neuropeptide in promoting sleep, *flp-24* overexpression caused a modest increase in RIS activity (Figure 5H). We conclude that ALA can act as an upstream activator of RIS and that *flp-24* neuropeptides promote RIS activation. Thus, a model emerges in which EGF activates both ALA and RIS, with ALA inducing behavioral quiescence that includes the promotion of RIS activation, which induces sleep bouts.

### ALA rather than RIS support survival after stress

It has been shown that ALA is required for survival following cellular stress, suggesting that sleep promotes stress resistance[41, 42]. Cellular stress-induced sleep requires both ALA and RIS, and these neurons can also inhibit behavioral activity independently of each other, suggesting that *C. elegans* possess at least two types of behavioral inhibition, sedation induced by ALA and sleep bouts induced by RIS. We hence tested whether the organism’s survival after a heat shock depends on the sedation conferred by ALA or on sleep bouts induced by RIS. We heat shocked wild-type worms, *ceh-17(-)* (to ablate ALA), *aptf-1(-)* (to ablate RIS), or *ceh-17(-)*;*aptf-1(-)* double mutants (to ablate both ALA and RIS) and followed their survival[41]. As in previous results, the *ceh-17(-)* mutation reduced survival by about one third. *aptf-1(-)* mutants worms were indistinguishable from wild-type animals, and *ceh-17(-);aptf-1(-)* double mutants lived for slightly less time than *ceh-17(-)* (Figure 6). These results suggest that ALA activation is protective against cellular stress by inhibiting behavior independently of RIS-induced sleep bouts. This suggests that cellular stress causes two types of behavioral quiescence that have different physiological functions. Sedation through ALA appears to be important in supporting survival after heat shock, and RIS induces sleep bouts whose functions are yet to be determined in the response to cellular stress.

**Figure 6.**
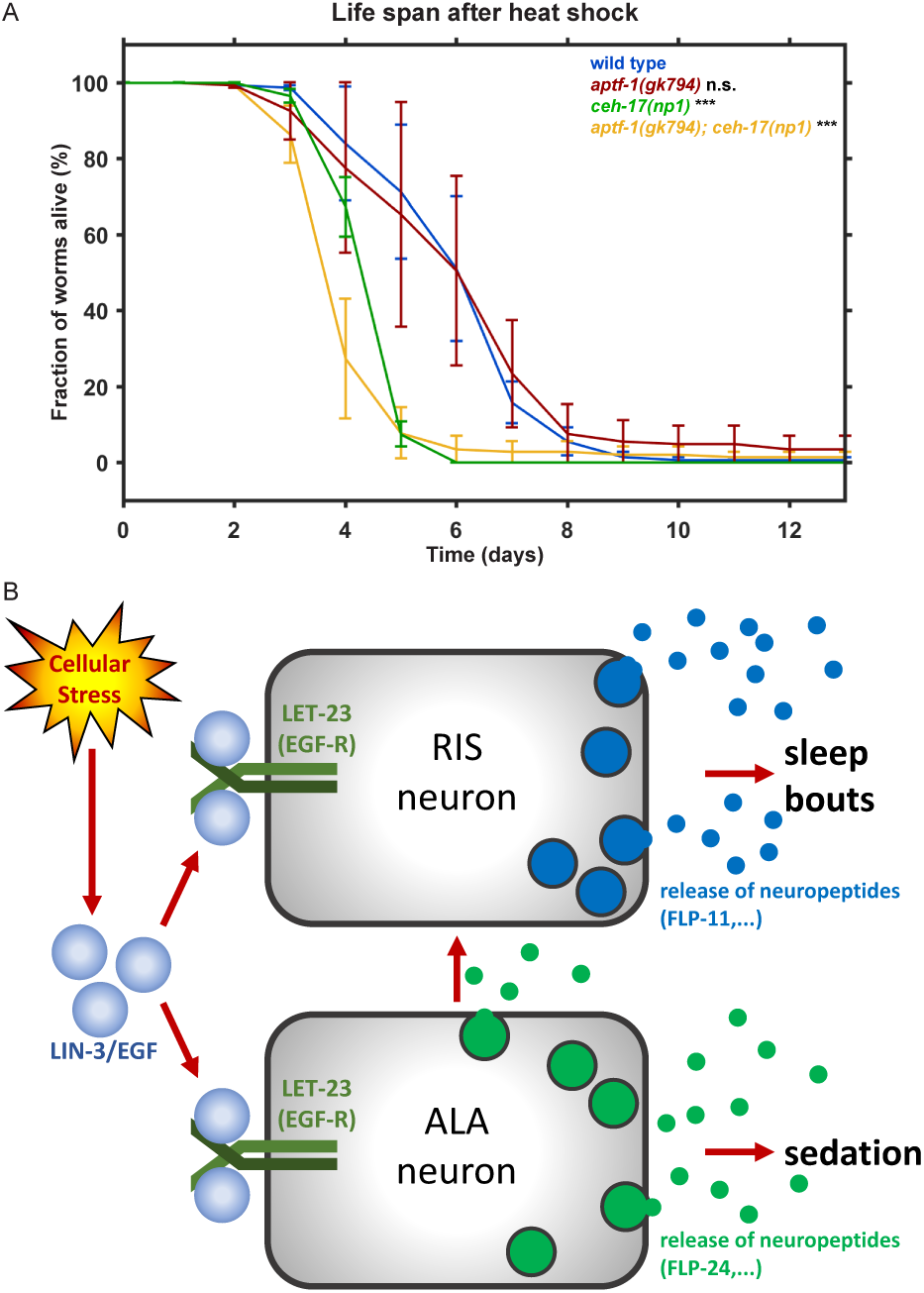
ALA (sedation) rather than RIS (sleep bouts) is required for survival following heat stress. (A) Survival of ALA and RIS mutants after heat shock (40°C, 20 min). *aptf-1lf* (red, deletion of RIS) did not show a different survival rate after the heat shock. *ceh-17lf* (green, deletion of ALA) and the *ceh-17lf*; *aptf-1lf* (yellow) double mutant died earlier than wild-type worms. Three biological replicates were performed, with N = 50 worms per allele per replicate. *** denotes statistical significance with p < 0.001, Cox proportional hazards regression. (B) Model for EGF-induced sedation and sleep through ALA and RIS neurons. Cellular stress leads to EGF release, which activates the RIS and ALA neurons via the EGF receptor. Both neurons release neuropeptides. ALA induces sedation and activates RIS. RIS induces sleep bouts.

## Discussion

### ALA is a sedating and sleep-promoting neuron

ALA is activated by EGFR and has been shown to act through multiple sleep-promoting neuropeptides, including those encoded by the *flp-13*, *nlp-8*, and *flp-24* neuropeptide genes. The peptides encoded by these genes act in parallel to induce sleep behavior and inhibit specific behaviors such as feeding[37–39]. Our data indicate that the role of ALA in sleep induction following EGF released can be best understood if it is viewed as a sedating and sleep-promoting neuron. According to this model, the neuropeptide cocktail released by ALA upon EGFR activation induces two different effects on the nervous system. First, cellular stress caused by heat shock inhibits behavioral activity such as feeding independently of RIS[40]. In this first ALA function, neuropeptides may inhibit wake-promoting neurons directly, and thus inhibit active behaviors causing sedation or lethargy over prolonged time [38, 39]. Sedation has some similarities to sleep, such as reduced voluntary movement and reduced responsiveness to stimulation but differs from sleep in that it does not display the fast switching properties that cause the succession of sleep bouts as well as a quick reversibility. Nevertheless, sedation is associated with increased sleep, indicating that these two behaviors are causally linked. The second mechanism acts through RIS, which induces sleep bouts.

### EGFR activates RIS to induce sleep bouts following cellular stress

EGFR activation in many tissues, including ALA, has been shown to act through PLC gamma[37]. Thus, EGFR most likely acts similarly on RIS by increasing its excitability through PLC signaling mechanisms. Increased excitability in turn could increase the probability of strong RIS calcium transients. The role of Ras signaling in RIS will need to be determined in future studies but speculatively could play a role in increased neurotransmitter expression. RIS activation allows fast neural dampening that affect brain and behavioral activity systemically[31, 33, 34]. RIS depolarization correlates with sleep bout induction during many conditions[31, 33]. Consistent with these previous findings, RIS activation also correlates with quiescence bouts following cellular stress. Several calcium transients follow rhythmically after cellular stress, with each calcium transient correlating with a sleep bout, indicating that RIS is sleep-active during stress-induced sleep. The fast-switching kinetics of RIS thus appear to account for the dynamic sleep bouts that occur following cellular stress. RIS also activated during the heat shock, consistent with a previous report [58], and functional RIS was partially required for immobilization, suggesting that immobility during the heat shock is promoted by RIS but that there is an additional mechanism that paralyzes it. In summary, EGFR acts through two parallel neurons, ALA and RIS, with ALA inhibiting specific behaviors on longer time scales and RIS inhibiting systemic behavior on shorter time scales. By engaging both ALA and RIS, EGFR signaling could thus inhibit behavioral and physiological activity with varying specificity and on different time scales.

### Sedation is protective after cellular stress rather than sleep bouts

Sleep is thought to play a protective role in coping with stress and sleep and sedation are induced by sickness. Across species, sleep is associated with increased vitality, but the relative contributions of sedation and sleep to recuperation from illness have not been disentangled. Loss of functional ALA has been shown to reduce viability following cellular stress [41]. We confirmed the essential role of ALA in surviving following heat shock. Interestingly, RIS was not required for survival under the conditions tested. This suggests that ALA becomes beneficial not through its role in RIS activation and sleep bouts, but through its RIS-independent sedating action. ALA has been shown to inhibit feeding independently of RIS[40]. Reduced food intake increases health and longevity [61], suggesting that ALA might, at least in part, act by reducing food intake following cellular stress. EGFR signaling has been shown to counteract aging and promote health in *C. elegans*. EGFR gain of function increases lifespan and promotes health, whereas loss of function of EGFR signaling increases the rate of aging[62, 63]. EGFR signaling has been shown to promote longevity by increasing the activity of the ubiquitin proteasome system during development in epithelia. This effect is mediated by Ras signaling and might cause greater fitness leading to a longer lifespan[64, 65]. Thus, EGFR can positively influence health and lifespan by acting at different points in developmental time through different mechanisms including transcriptional changes as well as changes in behavior. In summary, ALA and RIS play distinct functions in inhibiting behavior and sedation appears to play a major role in coping with stress. Likely, sedation and sleep also play distinct protective roles also in other organisms including humans.

## Acknowledgments

Some strains were provided by the CGC, which is funded by NIH Office of Research Infrastructure Programs (P40 OD010440). This work was supported by the Max Planck Society (Max Planck Research Group), and by a European Research Council Starting Grant (ID: 637860, SLEEPCONTROL) to HB. R. M.W. and D. M. M. were supported by R01 NS100547 and R01 NS081259 from the NIH and S. P. by a NSF Predoctoral Fellowship. This paper is dedicated to the memory of Max Fritz, who passed away unexpectedly in 2018 at the age of 20 years.

## Author Contributions

J.K. designed, performed, and analyzed the sleep experiments, drafted the methods part and edited the manuscript. M.F. identified the RIS transcriptome from sc-RNA-seq data. S.S. and A.H. generated the conditional *let-23* allele. W.S.C., A.O., and A.G. contributed to the generation of preliminary FACS RIS transcriptome data. R.M.W., S.P., and D.M.M. generated and analyzed the FACS RIS transcriptome. A.L. performed statistical comparisons of the transcriptomes. H.B. conceived the overall outline of the project, designed experiments, supervised the work, and wrote the manuscript. A.H., A.G., D.M.M., and H.B. acquired funding. All authors approved the final version of this manuscript.

## Declarations of Interest

The authors declare that they have no competing interests.

## STAR Methods

### CONTACT FOR REAGENT AND RESOURCE SHARING

Further information and requests for resources and reagents should be directed to the Lead Contact, Henrik Bringmann.

### EXPERIMENTAL MODEL AND SUBJECT DETAILS

#### Worm maintenance and strains used in this study

*C. elegans* was cultured on Nematode Growth Medium (NGM) agarose plates seeded with *E. coli* OP50 and incubated at 20°C [66, 67]. The following *C. elegans* strains were used:

| strain | genotype | backcrossed |
| --- | --- | --- |
| AH12 | *gap-1(ga133) X*. | 4x |
| AH4970 | *let-23(zh130[let-23::FRT::GFP::LoxP::FLAG::let-23]) II*. |  |
| AH5059 | *let-23(zh131[FRT::let-23::FRT::GFP::LoxP::FLAG::let-23]) II*. | 4x |
| AQ866 | *ser-4(ok512) III*. | 5x |
| BC16236 | *dpy-5(e907) I*; *sEx16236[rCesT17E9.2a::GFP + pCeh361]*. |  |
| CB30 | *sma-1(e30) V*. | 1x |
| CB6785 | *unc-119(ed3) III*; *eEx650[ilys-4p::GFP + unc-119(+)]*. | 2x |
| CG625 | *unc-103(n1213) pha-1(e2123) III*; *him-5(e1490) V*; *rgEx235[plc-3p::YFP + pha-1(+)]*. |  |
| DA1641 | *lin-15B&lin-15A(n765) X*; *adEx1641[C09B7.1::GFP + lin-15(+)]*. |  |
| FX01512 | *rabx-5(tm1512) III*. |  |
| FX01556 | *tiam-1(tm1556) I*. |  |
| FX02304 | *ssup-72(tm2304) II*. |  |
| FX02584 | *mct-6(tm2584) X*. |  |
| FX02735 | *T21F2.1(tm2735) X*. |  |
| FX03878 | *pkg-2(tm3878) IV*. |  |
| FX04341 | *F57A8.4(tm4341) V*. |  |
| FX04393 | *nlp-37(tm4393) X*. |  |
| FX04503 | *kin-15(tm4503) II*. |  |
| FX05049 | *R11F4.2(tm5049) II*. |  |
| FX05160 | *F38B6.6(tm5160) X.* |  |
| FX05990 | *F40A3.7(tm5990) V.* |  |
| FX06682 | *C49H3.3(tm6682) IV.* |  |
| FX17081 | *cyn-10(tm6703) II.* |  |
| FX18630 | *mrpl-19(tm4843) I.* |  |
| FX30291 | *unc-25(tm2727) III.* |  |
| HA328 | *lin-15B&lin-15A(n765) X; rtEx233[nlp-11p::GFP + lin-15(+)].* |  |
| HA329 | *lin-15B&lin-15A(n765) X; rtEx234[nlp-13p::GFP + lin-15(+)].* |  |
| HBR1261 | *goels288[flp-11p::mKate2::unc-54-3'utr, unc-119(+)].* | 2x |
| HBR1361 | *goels304[flp-11p::SL1-GCaMP3.35-SL2::mKate2-unc-54-3'UTR, unc-119(+)].* | 2x |
| HBR1777 | *goels384 [flp-11p::egl-1::SL2-mKate2-flp-11-3'UTR, unc-119(+)].* |  |
| HBR183 | *cca-1(ad1650) X.* |  |
| HBR1857 | *ceh-17(np1) I; goels304[flp-11p::SL1-GCaMP3.35-SL2::mKate2-unc-54-3'UTR, unc-119(+)].* |  |
| HBR2026 | *C50F7.6(gk744131) IV.* | 4x |
| HBR2040 | *ckr-1(ok2502) I.* | 2x |
| HBR2043 | *R09F10.1(ok3119) X.* | 1x |
| HBR2047 | *zig-2(ok696) X.* | 1x |
| HBR2066 | *R02F2.4(gk382575) III.* | 2x |
| HBR2067 | *F32D1.3(gk232755) V.* | 4x |
| HBR2068 | *ncs-6(gk138627) II.* | 2x |
| HBR2073 | *gei-1(gk170616) III.* | 2x |
| HBR2086 | *srx-2(gk242439) V.* | 2x |
| HBR2088 | *ceh-17(np1) I; aptf-1(gk794) II.* |  |
| HBR2107 | *ilys-4(gk402093) IV.* | 4x |
| HBR2108 | *srd-32(gk468949) V.* | 2x |
| HBR2193 | *goeEx727[flp-24p::ReaChR::mKate2-unc-54 3'UTR, unc-122p::dsRed]; goels304[flp-11p::SL1-GCaMP3.35-SL2::mKate2-unc-54-3'UTR, unc-119(+)].* |  |
| HBR2195 | *let-23(sa62) II; goels304[flp-11p::SL1-GCaMP3.35-SL2::mKate2-unc-54-3'UTR, unc-119(+)]; him-5(e1490) V; syls197[hs::LIN-3C(cDNA) + myo-2p::dsRed + pha-1(+)].* |  |
| HBR2205 | *syEx1286[hsp16-41p::flp-24, myo-2p::dsRed, KS+]; goels304[pflp-11::SL1-GCaMP3.35-SL2::mKate2-unc-54-3'UTR, unc-119(+)].* |  |
| HBR2238 | *bqSi542[unc-47p::FLP D5 + unc-119(+)]; let-23(zh131[FRT::let-23::FRT::GFP::LoxP::FLAG::let-23] II.* |  |
| HBR2239 | *bqSi542[unc-47p::FLP D5 + unc-119(+)]* | 2x |
| HBR2256 | *goeEx737[flp-24p::SL1-GCaMP3.35-SL2::SL2-mKate2::unc-54 3'UTR, unc-119(+)].* |  |
| HBR2257 | *goeEx737[flp-24p::SL1-GCaMP3.35-SL2::SL2-mKate2::unc-54 3'UTR, unc-119(+)]; him-5(e1490) V; syls197 [hs::LIN-3C(cDNA) + myo-2p::dsRed + pha-1(+)].* |  |
| HBR227 | *aptf-1(gk794) II.* | 10x |
| HBR507 | *flp-11(tm2706) X.* | 7x |
| HBR763 | *C10C6.7(goe4) IV.* |  |
| HT1757 | *unc-119(ed3) III; wwIs34[ins-24p::GFP + unc-119(+)].* |  |
| HT1768 | *unc-119(ed3) III; wwIs35[ins-27p::GFP + unc-119(+)].* |  |
| IB16 | *ceh-17(np1) I.* |  |
| JD596 | *avr-15(vu227) V.* | 2x |
| JT734 | *goa-1(sa734) I.* | 3x |
| KJ355 | *csq-1(jh109) X.* | 6x |
| LSC1264 | *sprr-1(ok3685) IV.* | 4x |
| LSC87 | *lstEx6[pdf-2p::GFP + myo-3p::mCherry]. (pdf-2 = nlp-37)* |  |
| MB5 | *lin-15(ts); him5(e1490); rtEx277[Pnlp-8::GFP+lin-15(+)].* |  |
| MT4433 | *ced-6(n1813) III.* |  |
| MU1255 | *nhr-67(tm2217) IV.* | 2x |
| N2 | *wild type* |  |
| NY2072 | *ynIs72[flp-1p::GFP].* |  |
| OH14368 | *pha-1(e1323) III; him-5(e1490) V; otEx6710[srd-32p::GFP + pha-1(+)].* |  |
| OH4836 | *otIs7[zig-2::GFP + rol-6(su1006)].* | 2x |
| PHX530 | *nlp-11(syb530) II.* |  |
| PHX700 | *ilys-4(syb700) IV.* |  |
| PS1839 | *let-23(sa62) II.* |  |
| PS2746 | *dpy-20(e1282) IV; syEx234[let-23::GFP + pBS + (pMH86) dpy-20(+)].* |  |
| PS4886 | *plc-3(tm1340) II.* |  |
| RB1001 | *cpna-3(ok922) IV.* |  |
| RB1136 | *R05G6.10(ok1159) IV.* |  |
| RB1151 | *cft-1(ok1180) V.* |  |
| RB1245 | *rga-1(ok204) II.* |  |
| RB1399 | *T01H8.2(ok340) I.* |  |
| RB1431 | *mps-2(ok1631) II.* |  |
| RB1468 | *dkf-2(ok1704) V.* |  |
| RB1512 | *Y57A10A.24(ok1803) II.* |  |
| RB1523 | *C24G7.1(ok1822) I.* |  |
| RB1911 | *ins-27(ok2474) I.* |  |
| RB2110 | *C39B10.1(ok2789) X.* |  |
| RB2269 | *flp-34(ok3071) V.* |  |
| RB2472 | *nep-26(ok3412) II.* |  |
| RB2627 | *srg-69(ok3686) II.* |  |
| RB5001 | *ZC434.9(ok5212) I.* |  |
| RB5002 | *srx-125(ok5810) V.* |  |
| RB669 | *wee-1.1(ok418) II.* |  |
| UP148 | *sem-5(cs15) X.* | 2x |
| VC1309 | *nlp-8(ok1799) I.* |  |
| VC1795 | *nrfl-1(ok2292) IV.* | 1x |
| VC1909 | *flp-1(ok2505) IV.* | 1x |
| VC20129 | *gem-1(gk298521) X.* |  |
| VC20159 | *F19C6.3(gk290941) X.* |  |
| VC20199 | *pms-2(gk247737) V.* |  |
| VC20231 | *rgef-1(gk243610) V.* |  |
| VC20263 | *glb-23(gk205062) IV.* |  |
| VC20307 | *T21D12.12(gk191670) IV.* |  |
| VC20312 | *R08B4.2(gk326023) X.* |  |
| VC20416 | *nhr-55(gk947607) V.* |  |
| VC20449 | *K02D3.1(gk299318) X.* |  |
| VC20489 | *nhr-128(gk960157) V.* |  |
| VC20519 | *F54H5.5(gk335875) II.* |  |
| VC20522 | *npr-29(gk337633) III.* |  |
| VC20554 | *F53F4.17(gk251960) V.* |  |
| VC20624 | *glb-32(gk360316) V.* |  |
| VC20630 | *T22F3.7(gk362826) V.* |  |
| VC20743 | *lmd-4(gk389517) V.* |  |
| VC226 | *ida-1(ok409) III.* |  |
| VC2565 | *frpr-3(ok3302) V.* |  |
| VC273 | *tag-89(ok514) IV.* | 1x |
| VC30075 | *T12B3.2(gk409439) IV.* |  |
| VC30104 | *tpa-1(gk414216) IV.* |  |
| VC30160 | *ile-1(gk427192) I.* |  |
| VC30245 | *B0244.10(gk446359) III.* |  |
| VC40013 | *frpr-3(gk240031) V.* |  |
| VC40057 | *dmsr-6(gk161951) II.* |  |
| VC40140 | *B0416.3(gk481746) X.* |  |
| VC40198 | *lgc-4(gk509234) X.* |  |
| VC40407 | *F56D5.6(gk622448) IV.* |  |
| VC40500 | *Y57G11C.36(gk961271) IV.* |  |
| VC40589 | *ZK1307.7(gk708119), sri-46(gk708025) II.* |  |
| VC40598 | *F57B10.4(gk712994) I.* |  |
| VC40613 | *frpr-16(gk722062) II.* |  |
| VC40626 | *tkr-1(gk729500) III.* |  |
| VC40630 | *Y51H7C.13(gk732116) II.* |  |
| VC40679 | *C25G6.4(gk759467) X.* |  |
| VC40734 | *nhr-194(gk784872) V.* |  |
| VC40893 | *Y116A8B.4(gk869095) IV.* |  |
| VC975 | *egas-2(ok1477) V.* |  |
| ZW477 | *bkip-1(zw10) II.* | 4x |

### METHOD DETAILS

#### Molecular biology and transgenic strain generation

All constructs were cloned using the MultiSite Gateway system (Invitrogen, Carlsbad, CA) with pCG150 (Addgene plasmid #17247), which contains *unc-119(+),* as the destination vector for LR reactions [68]. For verification, all constructs were Sanger sequenced. Genes encoding GCaMP3.35 and ReaChR were used that were codon-optimized for expression in *C. elegans*[69]. The following plasmids were generated and used in this study:

| construct name | construct structure |
| --- | --- |
| K351 | *flp-24p::SL1-GCaMP3.35-SL2::SL2-mKate2::unc-54 3'UTR, unc-119(+)* |
| K358 | *flp-24p::ReaChr::mKate2-unc-54 3'UTR, unc-119(+)* |

#### Transformation by DNA microinjection

DNA microinjection was used for the generation of transgenic *C. elegans* strains as described[70]. A young adult hermaphrodite was transferred using a ∼0.5 µl drop of Halocarbon oil 700 (Sigma) into a ∼2 µl drop of the same oil on an agar pad. To generate the agar pad before the start of the injections, a drop of 3% agarose in water was placed onto a glass slide, flattened with a glass slide, previously dried for one hour on a 95°C heating block. The worm was gently positioned with an eyelash to fix it on the agarose surface. Next, the glass slide with the fixed worm was placed onto a microinjection microscope setup, which consisted of an inverted microscope (Nikon, Ti-S), a micromanipulator (Eppendorf, Patchman) and an electrical microinjector (Eppendorf, FemtoJet). A microinjection needle (Eppendorf, Femtotips 2), pre-filled with DNA, was mounted on the microinjector. The needle was filled with DNA solution containing TE buffer, the target construct DNA, a co-injection marker DNA and was filled up with pCG150 DNA (Addgene plasmid #17247 [68]) to a final concentration of 100 ng/µL. As co-injection marker coel::RFP (*unc-122p::RFP*) was used, which expresses a red fluorophore in the coelomocytes (Addgene plasmid #8938 [71]). The construct was injected at the following concentrations, *goeEx727*: K358 10 ng/µl, coel::RFP 10 ng/µl, pCG150 80 ng/µl

The needle was inserted carefully into one arm of the gonad with the help of a micromanipulator, and DNA solution was injected with an injection pressure of 29.0 psi for an injection time of 0.4 seconds. Constant pressure was at 2.00 psi. The needle was retracted from the gonad and the worm was recovered with a 2 µl drop of M9. The worm was extracted from the liquid using a platinum wire pick and a drop of bacteria and transferred to a fresh NGM plate. After growing the worm at 20°C for 48h, F1 larvae were inspected with a fluorescence microscope for the expression of the co-injection marker and positive transformants were selected.

#### Transformation by microparticle bombardment

A second method used for the creation of transgenes was gold microparticle bombardment. *unc-119(ed3)* were used for bombardment and transformants were selected based on phenotypic rescue conferred by the *unc-119(+)* present in the plasmid that was used for transformation [72, 73]. Gold microparticles (chemPUR) sized 0.3-3 µm were coated with the DNA using spermidine (Sigma-Aldrich, 50 mM) and polyvinylpyrrolidone (Sigma-Aldrich, P-5288, Mol. 360, 0.1 mg/ml in 96% EtOH). Synchronized young adult worms were transferred onto NGM plates, which contained a 1 cm diameter bacterial lawn in their center and were cooled down by placing them on ice prior to transferring the worms. 20 µl of gold particle suspension was loaded onto the filter of a particle gun (Caenotec, Braunschweig). Helium (purity 5.0) was used at 8 bar to accelerate the particles into a vacuum chamber (−0.4 bar) onto the worms. Each construct was transformed eight times. Worms were recovered by cutting each NGM plate into six pieces after transformation and placing each piece onto a 12 cm NGM plate. Transformants were selected after two weeks incubation of the plates at 25°C. To select motile transformants, a 1 × 1 cm piece of an NGM plate seeded with OP50 was placed onto the plate and transformants were removed 0.5 −1 h later from the bacterial lawn.

#### CRISPR-based gene editing

##### Generation of the conditional *let-23* allele

An overview of the editing of the *let-23* locus can be found in Figure S4.

Plasmid constructs used:

pJe16: (single guide targeting *let-23* to insert *frt::gfp*)

Spacer sequence Sg3: TCAGCCTCTTCTGATGTTTC; in pDD162 (Addgene plasmid #47549) generated with OJE 299 and OJE 300.

pSS4: (single guide targeting *let-23* to insert *frt::gfp*)

Spacer sequence let-23::GFPsgRNA2: AAAACCTGAAACATCAGAAG; in pDD162 (Addgene plasmid #47549) generated with OSS92/ OSS73

pSS8: (single guide targeting *let-23* to insert 2nd *frt* site)

Spacer sequence sgRNA2_let-23BtyrK: TCAAGTGAATGCATACCCAT, in pDD162 (Addgene plasmid #47549) generated with OSS 84/ OSS85

pSS9: (single guide targeting *let-23* to insert 2nd *frt* site)

Spacer sequence sgRNA3_let-23BtyrK: TGGGTATGCATTCACTTGAA, in pDD162 (Addgene plasmid #47549) generated with OSS 86/ OSS87

pSS11: (precursor repair template targeting *let-23* to insert *frt::gfp*)

The pDD282 (Addgene #66823) plasmid was digested with the AvrII and SpeI restriction enzymes, to insert specific homology arms by means of the Gibson protocol[74]. The inserted homology arms were amplified by fusion PCR from genomic DNA with the following oligonucleotides:

3’Homology arm 1: OSS95/OSS96 (0.46 kb)

3’Homology arm 2, insertion of *frt* site: OSS97/OSS98 (0.14 kb)

3’Homology arm 1 and 2 were fused with: OSS 95/ OSS98 (0.6 kb)

5’Homology arm: OSS93/ OSS94 (0.81 kb)

pSS12: (*zh130* repair template)

pSS11 was used as template to amplify:

Fragment 1: OSS107/ OSS108 (5.7 kb)

Fragment 2: OSS106/ OSS26 (4.8 kb).

Fragments where then assembled by Gibson cloning[74]. This repair template integrates *gfp* as well one *frt* site 112 bp upstream of the linker sequence into the *let-23* locus.

OSS99: (zh131 repair oligonucleotide)

OSS99 (106 bp) was ordered as purified double-stranded oligonucleotide (Microsynth AG).

#### CRISPR/CAS9-generated alleles

AH4970 *zh130* [*let-23::FRT::GFP::LoxP::FLAG::let-23*] II.:

To insert the *frt::gfp::3xFlag* sequence in the 3’ region of the *let-23* locus, the CRISPR/Cas9 system according to Dickinson et al. (2015) was applied. The repair template plasmid pSS12 was injected into the N2 strain at a concentration of 10 ng/µl, the two single guides with integrated CAS9 plasmids pJE16 and pSS4 at a concentration of 25 ng/µl each together with the co-injection markers pGH8 (Addgene 19359) at 10 ng/µl, pCFJ104 (Addgene 19328) at 5 ng/µl and pCFJ90 (Addgene 19327) at 2.5 ng/µl. Further Selection was done according to Dickinson et al. (2015)[75].

AH5059 *zh131* [*let-23::FRT::let-23::FRT::GFP::LoxP::FLAG::let-23*] II.:

To insert the second *frt* site 2 kb upstream of *frt::gfp* and 5’to the protein kinase domain in the *let-23* locus, the repair oligonucleotide OSS99 was injected into *zh130* at a concentration of 500nM, the two single guide with integrated CAS9 plasmids pSS8 and pSS9 at a concentration of 25 ng/µl each with the Co-injection marker pCFJ90 (Addgene 19327) at 2.5 ng/µl. Co-injection marker was used to select for candidates which then were screened for positive insertion events by PCR using OSS118/ OLM137. *zh131* was sequenced to confirm the correct sequence and out crossed with N2 four times to get rid of any background mutations.

The following alleles and transgenes were used:

*bqSi506* [*rgef-1p::FLP D; unc-119*(+)] IV. [76]

*zh130* [*let-23::FRT::GFP::LoxP::FLAG::let-23*] II.

*zh131* [*let-23::FRT::let-23::FRT::GFP::LoxP::FLAG::let-23*] II.

The following oligonucleotides were used:

OJE 299:

CTATTGCGAGATGTCTTGTCAGCCTCTTCTGATGTTTCGTTTTAGAGCTAGAAATAGC AAG

OJE 300:

CTTGCTATTTCTAGCTCTAAAACGAAACATCAGAAGAGGCTGACAAGACATCTCGCA ATAG

OSS92: cttgAAAACCTGAAACATCAGAAG

OSS73: aaacCTTCTGATGTTTCAGGTTTT

OSS96:

GAAGTTCCTATACTTTCTAGAGAATAGGAACTTCTTTTAAAAGTAACTTTTTTAATGC G

OSS97: CTAGAAAGTATAGGAACTTCTAATAAATTTTCAGAGCTATCCCC

OSS98: GAGGCTCCCGATGCTCCaGGTTTTTTAGTTTTGAGATGTGG

OSS95: CGGCCAGTCGCCGGCACTAGGATGGGTCAACCGCACAGGAAG

OSS93: GGATGACGATGACAAGAGACTAGgAAACATCAGAAGAGGCTGAAGC

OSS94: GCTATGACCATTTATCGATTTCCTAGcctagtttaagctgcgccacaacag

OSS26: CTTGTAGAGCTCGTCCATTCCGTG

OSS106: ACAAGAGAgAAACATCAGAAGAGGCTG

OSS107: GATGTTTcTCTCTTGTCATCGTCATCC

OSS108: CACGGAATGGACGAGCTCTACAAG

OSS84: cttgTCAAGTGAATGCATACCCAT

OSS85: aaacATGGGTATGCATTCACTTGA

OSS86: cttgTGGGTATGCATTCACTTGAA

OSS87: aaacTTCAAGTGAATGCATACCCA

OSS118: GGCTATGGAGGATGTAAGCAGTGC

OLM137: CTATACTTTCTAGAGAATAGGAAC

OSS99:

GTAGAGTATTTTCTGAAAGAAACTTTAATTTCCAATTGAAGTTCCTATTCTCTAGAAAGTATAGGAACTTCGGTATGCATTCACTTGAATTGATGAATAATTGGAGGAATAATTG GAG

#### Generation of the *ilys-4* deletion

The *ilys-4(syb700)* deletion allele was generated by SunyBiotech using CRISPR/Cas9 and deletes 1270 nucleotides (the entire coding region) of the *ilys-4* gene between the following flanking sequences: gttcttgtgcgaatatctgaaattttattgtcattcagattcagattttcagatgtctacactggaaacgaatatcatgatcttacagcagacattctcattatcagatttgaaagaacataaattgcactgaaaaacatattaaaaattttgctccttccctatccaaaatatatataataaataaattagaaatttcatattttaaaattacagtcacagatgaagccgccgtgaatgaaaaaactcccaaa-tgaacattttttaatacttcagtcgcatatttatattgggaagagcattttctgtgccacattgctctctaataaataattattttactatacggagattttcgagctactgaactaaattgaa

#### Strain generation by genetic crossing

Transgenes were backcrossed two times against N2 to remove the *unc-119(ed3)* mutation. For following genotypes during strain crossing, the animals were genotyped using either Duplex PCR genotyping of single worms (mostly to detect deletions) [77], tetra-primer ARMS-PCR (to detect single nucleotide changes)[78], or sanger sequencing (to detect single nucleotide changes). Primers used for genotyping are listed in Table S5.

#### Transcriptome extraction from single-cell RNA sequencing data

For single-cell sequencing, the data set from Cao *et al.* was used in this analysis[43]. To identify the transcriptome cluster corresponding to RIS within the neuronal sci-RNA-seq clusters we used our previous observations that only RIS strongly and specifically expresses *flp-11* neuropeptides [32]. Gene counts and t-SNE based cell clusters were used as provided by the authors. Using the expression of the marker gene *flp-11*, one cluster was identified as the RIS cells. Cells with less than 70 UMI counts were discarded from the analysis. Only genes with at least one count in at least 5 cells were considered in the subsequent analysis. Differential expression analysis was done using edgeR (version 3.24.3) [79], fitting a negative binomial generalized log-linear model to the read counts for each gene. P values are results from a likelihood ratio test and were adjusted for multiple testing using Benjamini-Hochberg. Differential expression analysis was performed twice, once comparing RIS genes to all other remaining genes and once comparing RIS genes to all other cells. The significance level was set to alpha = 10% for all statistical tests. All analyses have been performed in R (version 3.4.0; R Core Team 2018). The single cell count data by Cao *et al.* contains counts for 20271 genes in 42035 cells derived from L2 larvae[43]. Cluster 13 was identified as the RIS cell cluster, containing 45 RIS cells.

#### Differential expression analysis of RIS versus all neurons

Here the analysis was conducted on the 7603 neuronal cells only. Post filtering, there were 9497 genes available in 7448 cells (of which 44 were RIS cells) for differential expression analysis. 8100 genes were down regulated in RIS according to this analysis of which 6 were statistically significant. 1331 genes were up regulated of which 60 were statistically significant. The most strongly enriched gene was *flp-11*, with an enrichment of 157-fold. Differential genes listed in Table S1.

Comparing the differentially and significantly expressed genes from the single-cell sequencing data set with the differentially and significantly genes from the bulk sequencing data set there were 58 genes present in both data sets. Comparing all differentially expressed genes from the single-cell sequencing data set with the differentially and significantly genes from the bulk sequencing data set there were 479 genes present in both data sets.

#### Differential expression analysis of RIS versus all cells

Here the analysis was conducted on all 42035 cells from the single-cell data set[43]. Post filtering, there were 9497 genes available in 39634 cells (of which 44 were RIS cells) for differential expression analysis. The results were compared to the results obtain from Bulk-RNAseq data. 7719 genes were down regulated in RIS according to this analysis of which 138 were statistically significant. 1410 genes were up regulated of which 243 were statistically significant. The most strongly enriched gene was *flp-11*, with an enrichment of 588-fold. Differential genes listed in Table S2.

Comparing the differentially and significantly expressed genes from the single-cell sequencing data set with the differentially and significantly genes from the bulk sequencing data set there were 228 genes present in both data sets. Comparing all differentially expressed genes from the single-cell sequencing data set with the differentially and significantly genes from the bulk sequencing data set there were 691 genes present in both data sets.

#### Bulk sequencing of FACS-isolated cells

RIS neurons, marked with mKate2 were isolated by FACS from L2 larval animals of genotype HBR1261[*goels288(flp-11p::mKate2::unc-54 3’UTR + unc-119(+))*] [32]. Cell dissociation and FACS methods were as previously described [45]. Briefly, synchronized L1 larvae were plated on NGM media seeded with OP50-1 bacteria and grown overnight at 23°C. L2 stage larvae were dissociated by successive treatments with 0.25 % SDS, 0.2 M DTT and 15 mg/ml pronase. A 5 µm filter was used to remove large debris and DAPI was added to mark dead cells. mKate2-labeled RIS neurons (8,000 – 50,000 cells/sample) were isolated by FACS (BD FACS Aria) and collected in Trizol for RNA extraction. Reference RNA samples were obtained from quick frozen aliquots of synchronized L2 larvae obtained immediately before initiating the cell dissociation protocol. The Clonetech-Takara SMART-Seq V3 Ultra Low Input RNA Kit was used for cDNA synthesis and amplification. Paired-end-100 (PE-100) data were collected in an Illumina NovaSeq 6000. ≥ 60 million reads/sample were collected for three independently isolated RIS samples and for three L2 whole animal reference samples.

To analyze data for bulk sequencing, Quality Control of the input reads was done using fastQC (version v0.11.2; Andrews, Simon, 2014, “FastQC A Quality Control tool for High Throughput Sequence Data” https://github.com/s-andrews/FastQC). Star (version 2.4.0) was used to align reads to the reference assembly WBcel235 of *Caenorhabditis elegans* [80]. Gene annotation was used from release 94. Multiqc (version 1.5) was used to facilitate quality control on the input data as well as the alignment statistics [81]. Gene level counts were generated using RSEM (version 1.2.19) to deal with multimapping reads [82]. All downstream analyses have been performed in R (version 3.4.0; Core Team, 2018, *R: A Language and Environment for Statistical Computing*. Vienna, Austria: R Foundation for Statistical Computing. https://www.R-project.org/). Read counts were normalized using tximport (version 1.8.0) [83]. CPM values were generated for first unbiased analyses. Correlation based clustering and a PCA analysis were conducted to assess sample structure and identify potentially problematic samples. Differential expression analysis was done using edgeR (version 3.24.3) fitting a negative binomial generalized log-linear model to the read counts for each gene [79]. P values are results from a likelihood ratio test and have been adjusted for multiple testing using Benjamini-Hochberg. The significance level was set to alpha = 5% for all statistical tests.

Three biological replicates of isolated RIS and three biological replicates of control cells (all cells) were collected and bulk sequenced. One RIS sample was excluded from the analysis as it did not cluster with the other replicates. 4504 genes were down regulated in RIS according to this analysis of which 3183 were statistically significant. 3197 genes were up regulated of which 1188 were statistically significant. Among the four most strongly enriched genes was *flp-11*, with an enrichment of 890-fold. Differential genes are listed in Table S3.

#### Differential expression analysis of ALA and comparison with RIS

Genes expressed in ALA were also extracted from the data set from Cao *et al.* as above[43]. To identify the transcriptomes corresponding to ALA we used the previous observations that ALA expresses *flp-24, flp-13,* and *flp-7* neuropeptides [38, 39]. Cells with less than 70 UMI counts were discarded from the analysis. Only genes with at least one count in at least 5 cells were considered in the subsequent analysis. Here the analysis was conducted on the 7603 neuronal cells only. Post filtering, there were 9497 genes available in 7448 cells for differential expression analysis. 22 cells, which formed part of cluster 11, were identified as ALA[43]. Differential expression analysis was done using edgeR [version 3.24.3; @edgeR] fitting a negative binomial generalized log-linear model to the read counts for each gene [79]. P values are results from a likelihood ratio test and have been adjusted for multiple testing using Benjamini-Hochberg. The significance level was set to alpha = 10% for all statistical tests. Differential expression analysis was performed comparing ALA cells to the remaining pan-neuronal cells. 8286 genes were down regulated in RIS according to this analysis of which 0 were statistically significant. 1189 genes were up regulated of which 22 were statistically significant. Among the top enriched genes were *flp-24*, *let-23*, *flp-7*, and *nlp-8*, which have previously been demonstrated to be expressed in ALA, indicating that the ALA transcriptome was correctly identified [37, 39]. Differential genes listed in Table S4. Pairwise correlations of logFC from tests vs pan-neuronal background were computed. Columns and rows were ordered following hierarchical clustering. All neuronal clusters with less than 100 cells were compared to the remaining pan-neuronal background. Based on the resulting logFC, pairwise correlations and hierarchical clustering were calculated.

#### Long-term imaging using hydrogel microchambers

Imaging of behavior and calcium activity was performed using Agarose Microchamber Imaging (AMI) as described [84, 85]. Shortly, a polydimethylsiloxane (PDMS) mold was used to create microcompartments from melted 3% high-melting agarose (Fisher Scientific GmbH) dissolved in S-Basal[67]. The following chamber sizes were used: 190 µm × 190 µm × 15 µm (X length x Y length x Z depth) for L1, 370 µm × 370 µm × 45 µm for adults. The microchambers were filled with either eggs (for L1 arrest experiments) or young adults (for heat shock experiments), sealed with a cover slip, and attached with double-side adhesive tape (Sellotape) into an opening milled into a 3.5 cm plastic Petri dish. An additional 2 ml volume of 3 % high melting agarose was filled to form a ring around the agar block containing the microcompartments, serving as a moisture reservoir. The space between the agarose pad and the agarose ring of the Petri dish was filled with melted 3% low melting agarose dissolved in S-Basal. The sample equilibrated for at least 2 h before the start of imaging. For imaging, a home-made heating lid was used that kept the temperature at 25°C to avoid condensation.

#### Microscopy setups for imaging

Imaging was performed on either a TiE or Ti2 inverted microscope (Nikon) with an automated XY stage (Prior, Nikon). The following objectives were used: 40x 0.45 NA dry, or 60x 1.4 NA oil for reporter co-expression experiments, 10x NA 0.45 dry with DIC filter for L1 arrest experiments and 20x NA 0.75 dry with an additional 0.7 lens placed in the c-mount of the camera for all experiments with young adult worms. Adults were imaged using the 10x objective. L1s were imaged with the 20x objective. This constellation allowed fitting 1 and 36 chambers simultaneously onto the camera chip for adults and L1, respectively. Microscopes were equipped with red-light (Semrock BrightLine HC 785/62, 45 mm diameter) dia illumination for differential interference contrast (DIC), which was used for behavioral imaging. Standard filter sets were used for GFP/GCaMP (ET-EGFP, Chroma) and mKate2 (TexasRed, Chroma) fluorescence imaging and optogentic stimulation. Images were acquired using either am electron multiplying charge-coupled device (EMCCD) camera (iXon DU-897D-C00-#BV, 512 × 512 pixels, Andor) or back-illuminated sCMOS camera (Prime 95B, 1174 × 1174 pixels, Photometrics) for fluorescence imaging. For experiments requiring only DIC imaging, an sCMOS camera (Neo, 2560 × 2160 pixels, Andor) was used. For fluorescence illumination and optogenetics an LED system was used (CoolLED). The LED system provided light with the wavelength of 488 nm for GFP excitation and 565 nm for mKate2 excitation and was triggered via the transistor-transistor logic (TTL) “fire out” signal of the camera. The software used to control the microscope and image acquisition was either iQ2/iQ3 (Andor) or NIS elements (Nikon).

#### Calcium imaging and optogenetics

For 490nm illumination for GCaMP imaging, light intensity was 0.16 mW/mm^2^ using a 20x objective. EM gain was set to 200 and exposure time was 20 ms. For 565 nm illumination (mKate2 imaging), light intensity was 0.06 mW/mm^2^ using a 20x objective. Light intensities were quantified using a light voltmeter (PM100A, Thorlabs). Samples were fixed on the microscope for long-term imaging experiments using a home-made aluminum sample holder for 3.5 cm plastic dishes. For ReaChR experiments, worms were fed with all-*trans* Retinal (Sigma, ATR). 20 µL of a 0.2 mM all-*trans* Retinal solution was added to a seeded NGM plate and L3/L4 worms were placed on it. The plate containing the worms was stored in the dark at 20°C in an incubator and the worms were used for optogenetic experiments the following day. For control experiments worms grown without ATR were used.

For optogenetic experiments worms were placed into microchambers and imaged at a frame rate of 0.3 frame/s. The optogenetic experiment consisted of three parts. First, RIS GCaMP baseline activity was recorded for 5 min, followed by a 5 min optogenetic activation period (1.09 mW/mm^2^) while we continued to record GCaMP fluorescence. After the end of the activation period an additional 5 min of GCaMP fluorescence was recorded. Green light illumination for optogenetic activation was shuttered so that it only occurred in between the acquisitions. Each worm was probed optogenetically for 3 to 4 times with a break of at least 2 hours in between each trial. All trials for each worm were averaged to obtain one N. Individual worms that did not express ReaChR in the ALA neuron were identified post hoc and were censored.

#### Reporter gene expression in RIS

Genes enriched in the RIS transcriptome were tested with existing reporter strains reported in the literature to be expressed in RIS. Reporter strains expressing GFP were crossed with an mKate2-expressing reporter strain for RIS. *mKate2* expression was driven via the *flp-11* promoter. Cross progeny animals were immobilized in a 5 µL drop of levamisole on a 200 µL high-melting agarose pad on a glass slide and covered with a cover slip. Co-expression of both fluorescent gene reporters was either tested with a spinning disc system (488 nm, 565 nm lasers, Andor Revolution, Yokogawa CSU-X1, Nikon TiE) or on a standard widefield fluorescence microscope setup (Nikon TiE, LEDs 488 nm, 565 nm). On both setups either 40x, 60x or 100x oil objectives were used. A z-stack was taken through the worm’s head and the maximum projection was calculated. The gamma values for each color channel were adjusted for display.

#### Mutant sleep screen during L1 arrest

L1 arrest screening was done with AMI. Usually five strains plus a wild-type (N2) control were filmed in one experiment. For this experiment, 12 pretzel stage eggs per strain were taken from a growing population and transferred into microchambers (190 µm × 190 µm x 15 µm). Each egg was transferred using an eyelash into an individual chamber while care was taken not to transfer any food. The eggs of each strain were arranged in adjacent microchambers so that they formed a characteristic pattern and thus were unambiguously identifiable. After the agarose microchambers were sealed, they were placed into an incubator at 20°C in the absence of light for 48 hours, during which time the worms hatched and arrested at the L1 larval stage. Then the arrested worms were imaged using DIC for 12 h with a frame rate of 0.2 frames/s and exposure time of 20 ms using a 10x objective combined with an additional 1.5x lens (total magnification was 150 x). Sleep bouts were extracted for individual worms using frame subtraction and mutants with either significantly decreased or increased sleep fraction were retested. If the mutant strain had not yet been outcrossed against N2 after mutagenesis, it was outcrossed two times before retesting. If the phenotype persisted, it was outcrossed for further two times (to a total of 4 x) and tested again. Only mutations that produced a significant sleep phenotype after 4 x outcrossing were scored as screen hits. 16 alleles were excluded from the screen as the worms showed severe developmental problems. Embryos from this strain often died during embryonic development or as an early L1 larvae. Survivors were small and had developmental defects preventing quantification of sleep. The excluded alleles were *sma-1(e30)*, *T21D12.12(gk191670)*, *glb-23(gk205062)*, *F54H5.5(gk335875)*, *glb-32(gk360316)*, *lmd-4(gk389517)*, *B0416.3(gk481746)*, *lgc-4(gk509234)*, *F57B10.4(gk712994)*, *frpr-16(gk722062)*, *Y116A8B.4(gk869095)*, *Y57G11C.36(gk961271)*, *nhr-194(gk784872)*, *ida-1(ok409)*, *nlp-8(ok1799)*, and *C39B10.1(ok2789)*.

#### Induction of cellular stress by heat shock

All heat shock experiments were performed in young adult worms before the first egg was laid. AMI was used with chambers of the size 370 µm × 370 µm × 45 µm. 8 to 10 young adult worms were transferred into a 5 µL drop of sterile distilled water placed on the agarose pad containing the microchambers with as little food as possible. While the liquid soaked into the agarose, individual worms were distributed into individual agarose microchambers with an eyelash. The microchambers were sealed with a cover slip and attached with double-faced adhesive tape to an opening of a metal plate that was part of a home-made temperature control device. The temperature control device contained the sample in a 10 × 10 mm opening of a metal plate (490 × 200 mm) and contact between the metal plate and the microchambers was created by filling the space with additional liquid agarose. The temperature of the metal plate and sample was measured by a Pt1000 temperature sensor that was placed in close proximity of the sample. Temperature was controlled by a Peltier element and its controller (Peltier-Controller TC0806, CoolTronic). The Peltier element transported energy from or to a metal grid acting as a heat sink, which itself was equilibrated with the surrounding air temperature using a small fan (Figure S3).

For the heat shock experiments, the device with the agarose pad and worms was stored in a dark 20°C incubator to equilibrate for 90 minutes. The device was then placed into a standard glass slide holder on an imaging microscope, connected to the Peltier controller and the temperature was set to 22°C. The plastic dish containing the microchambers was closed by a heated lid, whose temperature was set to 25°C to avoid drying out of the sample and condensation on the lid. Each worm was imaged for 3 hours with a sampling rate of 0.05 frame/s. In the first 60 minutes, baseline activity was imaged. Then the heating lid temperature was turned to 37.5°C and after 3 min the Peltier-Controlled metal plate was set to 37.0°C for a duration of 20 min to deliver the heat shock. To end the heat shock, the Peltier-Controller was set to 22°C and the lid was set to 25.0°C again. After the end of the heat shock, imaging was continued for an additional 2 h.

#### Induction of protein overexpression through temperature increase and *hsp-16.41p*

For overexpression of *lin-3* and *flp-24*, the *hsp-16.41* promoter and a temperature increase was used[37, 39]. The handling procedure of delivering this temperature increase for inducing gene expression was the same as the procedure of delivering a heat shock. The only differences were the length and the magnitude of the temperature stimulus. The length was slightly increased from 20 to 30 min but the temperature was increased to only 30.0°C and the heating lid to only 30.5°C, both for 30 min. Worms were filmed for another 6 hours after the temperature increase, at 22°C and with the lid set to 25.0°C. Control experiments without the heat shock-inducible transgene showed that this milder temperature increase was insufficient to trigger measurable stress-induced sleep.

#### Lifespan assay

Lifespan measurement were performed after heat shock similar to previously described. Briefly, a synchronized population of young adult worms was subjected to a heath shock and survival was followed [41, 86]. Worm populations were synchronized by isolating embryos and hatching them in the absence of food [87]. For each strain, two 6 cm plates full with gravid hermaphrodites were taken. Worms were harvested by washing them off with 2 ml M9, and transfer into a 1.5 mL Eppendorf tube. Worm were pelleted by centrifugation at 4.8 × 10^3^ rcf, the supernatant was removed and 500 µL of freshly prepared bleach solution was added to the pellet. To prepare the bleach solution, a stock solution of 1:1 1M NaOH solution and hypochlorite solution was diluted 1:2 with distilled water. Tubes with worms and bleach solution were mixed for 90 seconds by gentle manual agitation. Eggs were pelleted by centrifugation and the pellet was washed with 1 mL M9. Pelleting and bleaching was repeated and followed by three washing steps with 1 ml of M9 each.

The isolated eggs were resuspended in 1 mL M9 and transferred to a clean 1.5 mL Eppendorf tube. The tube was placed on a spinning shaker overnight. Next day the eggs had hatched and larvae were arrested at the L1 stage. 200 µL of each strain was pipetted on an NGM plate containing bacterial food. Worms were allowed to develop until the young adult stage in a dark 20°C incubator. For the heat shock a water bath (GFL, 1083) was heated to 40°C, and the correct temperature was verified by the internal and an additional external thermometer of the water bath (Greisinger electronic, GMH3710). The temperature was monitored during the whole heat shock process. For each strain 50 young adult worms were transferred onto 5 NGM plates, to obtain exactly 10 worms per plate. The plates were either seeded or unseeded with bacteria to do the heat shock either in the presence or absence of food. For the heat shock without food, L4 worms were picked with an eyelash to unseeded plates 12 hours before the heat shock. 24 hours after the heat shock all worms were picked to seeded plates again. Worms were starved for 36 hours in total. In the food condition, worms stayed on seeded plates all the time. These plates were sealed with parafilm and simultaneously placed into the water bath. The plates were placed into the water so that the half that contained the agar with the worms was down and submerged in the water. After 20 min, all plates were removed simultaneously from the bath and placed on ice for exactly 2 minutes. Water on the outside of the plates was removed with paper towels and the plates were stored in a dark incubator at 20°C. Every 24 h worm survival was counted by an experimenter that was bind to the genotype of the worms. Each worm that was not spontaneously moving was stimulated with a short pulse (10-20 s) of blue LED light delivered by a stereomicroscope (Leica, M165 FC). If the worm reacted to this light stimulus it was scored as “alive”. If no reaction was observed it was counted as “dead” and removed from the plate. Worms which could not be found on the plate, e.g. crawling up the plate wall and dry out, were counted as “censored”.

### QUANTIFICATION AND STATISTICAL ANALYSIS

#### Sleep detection using frame subtraction

All imaging data was saved as single .tif files and were further analyzed using home-made MATLAB (MathWorks) routines. Sleep bouts were defined by immobility, which was detected using a frame subtraction algorithm as described [50]. For the analysis, the image was cropped to only contain one microchamber containing one individual worm. For each frame, intensity values of each pixel were subtracted from the consecutive frame and the average of the absolute values for each frame was computed. The mean intensity was smoothed over 40 frames. The smooth function used was a robust version of a linear regression, which used weighted linear least squares and a 2nd degree polynomial model, by assigning lower weight to outliers in the regression (smooth(y,method,‘rloess’)). Intensities of the smoothed data were then normalized with 1 presenting the highest intensity value, and 0 the lowest intensity value. A sleep bout during L1 arrest was defined as a smoothed normalized value that was lower than 40% of the maximum intensity for at least 120 seconds. The sleep bouts extracted from the data set for each worm were used to calculate the mean sleep bout length, sleep bout frequency, and fraction of time spent in sleep bouts. Individual traces in which no sleep bouts were visible by manual inspection were scored as not containing any sleep bouts. The fraction of time spent in quiescence was used as a main criterion to score phenotypes in the genetic screen. Data for different individuals was averaged and statistically compared with wild-type N2 data obtained from an internal control (worms analyzed on the same agarose chip). In adult worms, sleep was defined by the same criteria as in L1 arrested larvae. To statistically compare after the heat shock, sleep data was binned by averaging data corresponding to time intervals of each 30 min following the heat shock.

Neuronal activity of the worms was measured with the green fluorescent calcium sensor GCaMP3 expressed from the RIS-specific *flp-11* promoter [32, 33, 88]. RIS was extracted based on fluorescence intensity using a home-made MATLAB routine. For RIS extraction, the pixels of each frame were binned 4:1 and the highest intensity pixel was identified that defined the center of the RIS neuron. The x-y position of this highest pixel was used to center a region of interest containing RIS and to crop this region from the original frame. The size of the region of interest was chosen to contain RIS and a limited amount of background (13 × 13 pixels for overexpression experiments and 21×21 pixels for optogenetic experiments). To identify RIS within the region of interest, its mean intensity was calculated and pixels that had a higher intensity than 25% of the mean of all pixels were counted as “signal”. Pixels below 25% mean signal intensity were counted as “background”. To calculate RIS intensity, the mean of all “background” intensities was subtracted from the mean of all “signal” intensities. Accurate tracking by the software algorithm was manually controlled at four time points (first frame, the frame after 1/3 of the movie, the frame after 2/3 of the movie, and the last frame in the movie). Image series in which RIS could not be identified automatically were censored. ALA position was identified by manually selecting the center for cropping a region of interest. For this procedure, a semi-automatic MATLAB routine was used that performed the same downstream data analysis as the automatically tracking routine.

Neural intensities measured before applying the heat shock were used as baseline and data was normalized as difference over baseline (ΔF/F). To determine sleep bouts in calcium imaging data sets, movement of the animal was detected based on the position of the center of the tracked head neuron. To extract sleep bouts, first the speeds were normalized, similar as described before but without any smoothing. Sleep was defined as time periods of less than 1.5% of the normalized movement speed.

#### Statistical tests

Statistical tests used were Wilcoxon rank tests for paired samples and Cox proportional hazards regression to test survival rates (both calculated in MATLAB). P values for differential expressed genes in the transcriptomes are results from a likelihood ratio test and have been adjusted for multiple testing using the Benjamini-Hochberg Procedure with a false discovery rate of 5% for FACS/RNA-seq data or 10% for sci-RNA-seq data, respectively (calculated with R). The specific tests used are described in the figure captions and the results section. The graphs show mean ± SEM unless noted. Compact boxplots were used for the visualization of L1 arrest screen data, with the box representing the 25%-75% range, the black dot representing the median and empty circles representing outliers. All other boxplots show individual data points, the box represents the 25%-75% range, and the thin gray line is the median. Whiskers for both types of boxplots correspond to approximately +/–2.7σ, which is 99.3% coverage if the data are normally distributed. Both types of boxplots were plotted via the “boxplot” function of MATLAB. For each experiment at least two biological replicates were performed and the number of biological replicates is stated in the figure legend.

## Supplementary information

**Table S1** sc-RNA-seq RIS transcriptome compared with all neurons

**Table S2** sc-RNA-seq RIS transcriptome compared with all cells

**Table S3** Bulk-sequenced FACS RIS transcriptome compared with all cells

**Table S4** sc-RNA-seq ALA transcriptome compared with all neurons

**Table S5** List of primers used for genotyping

**Figure S1.**
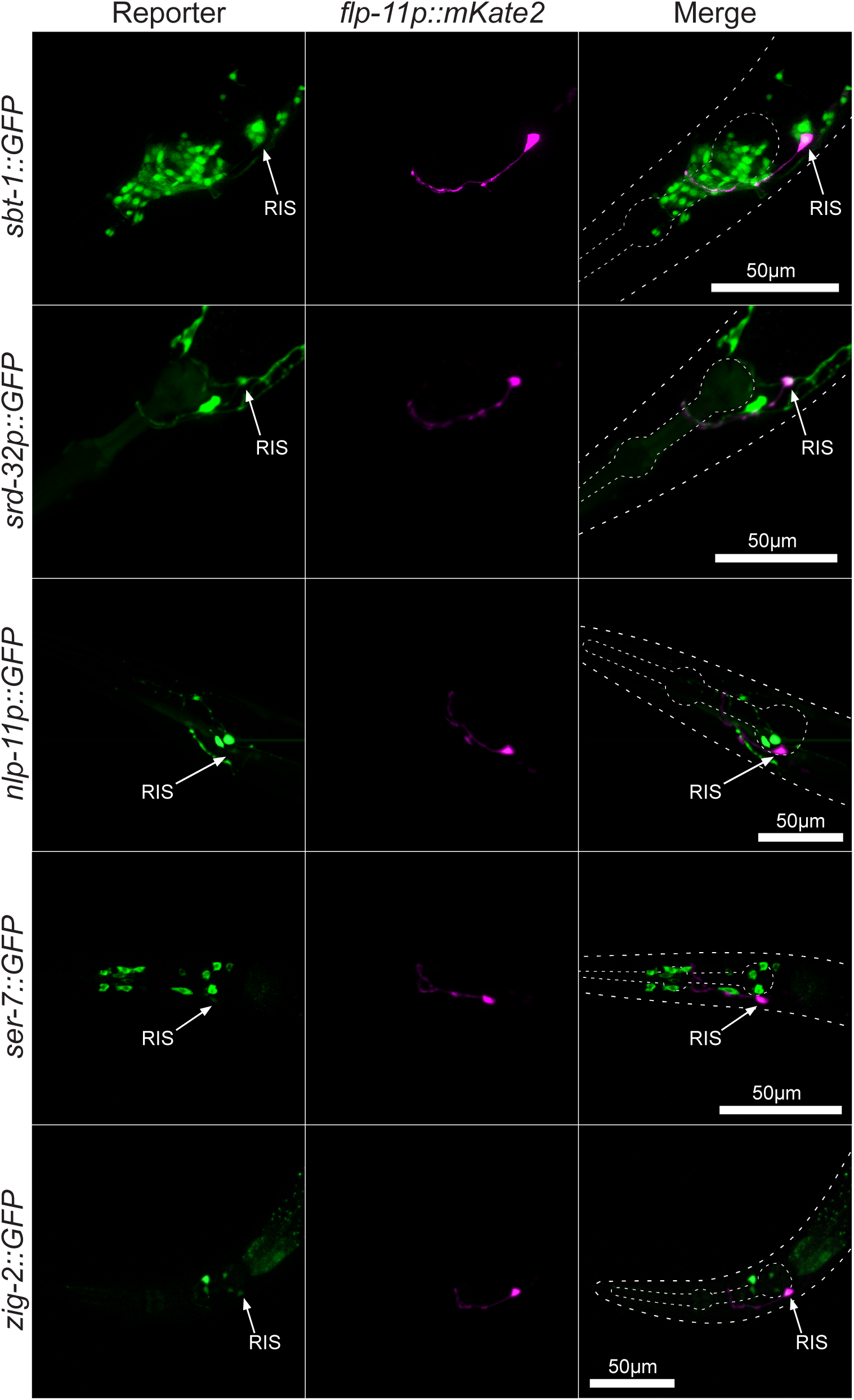
RIS-enriched genes for which fluorescence transgene reporters are expressed in RIS. Validation of RIS enriched genes using fluorescent transgene reporters. Example micrographs for *srd-32p::GFP*, *sbt-1::GFP*, *nlp-11p::GFP*, *ser-7::GFP*, *zig-2::GFP*, and their co-localization with *flp-11p::mKate2*. Dashed lines display the outlines of the head and pharynx (anterior is left, dorsal is up). RIS is indicated with a white arrow. Scale bar is 50µm.

**Figure S2.**
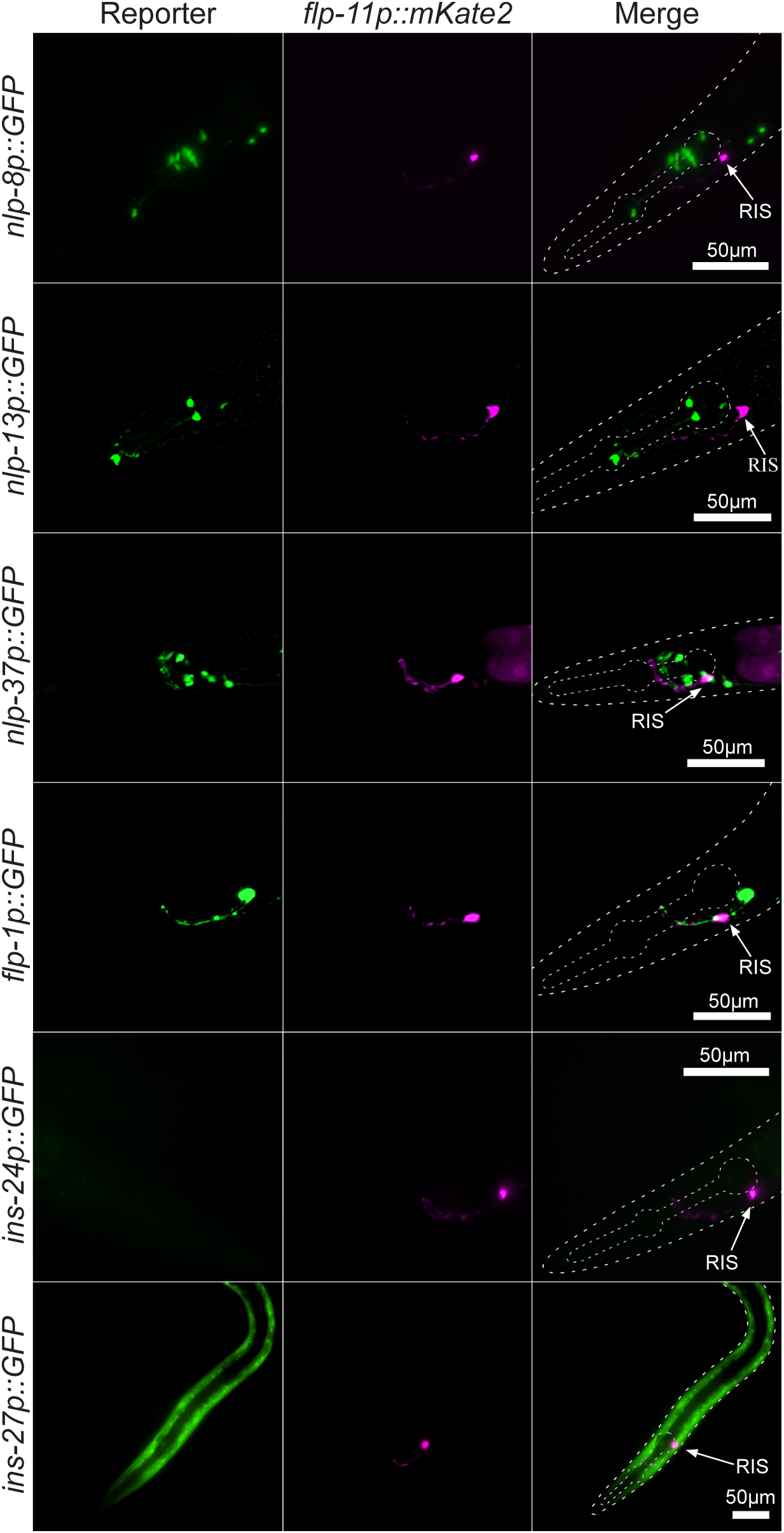
RIS-enriched genes for which fluorescent transgene reporters do not show RIS expression. Example micrographs for *nlp-8p::GFP*, *nlp-13p::GFP*, *nlp-37p::GFP*, *flp-1p::GFP*, *ins-24p::GFP*, *ins-27p::GFP*, and their co-localization with flp-11p::mKate2. Dashed lines display the outlines of the head and pharynx (anterior is left, dorsal is up). RIS is indicated with white arrows. Scale bar is 50µm. Lack of reporter gene expression could reflect false positives transcriptome results or false negative reporter expression.

**Figure S3.**
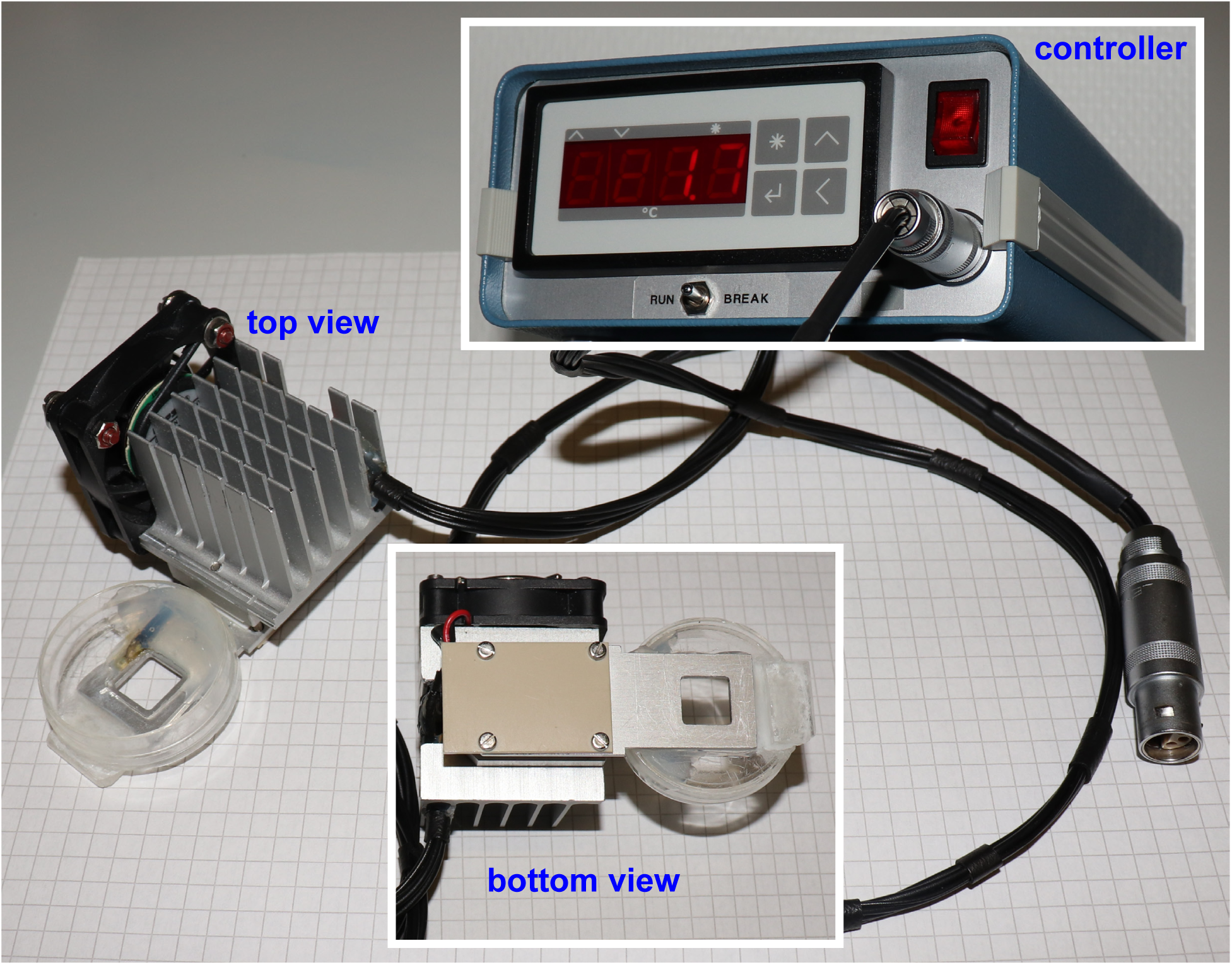
Heat control device. An agarose microchamber on a glass slide can be placed in the hole of the metal plate (bottom view, right side), which is attached to a Peltier element. Heat is transported by the Peltier element from the metal plate to a metal grid, which is equilibrated with the surrounding air temperature using a small fan (top view). A small petri dish is also glued to the metal plate to allow the filling with agarose, serving as a moisture reservoir and creating contact between the metal plate and the microchambers.

**Figure S4.**
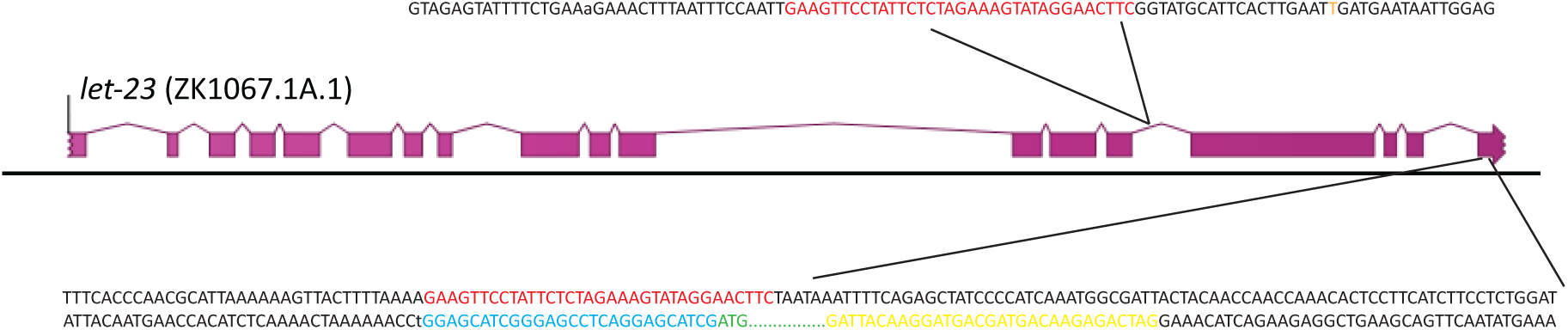
CRISPR/Cas9-based editing of the *let-23* locus. A schematic representation of the generation of the conditional *let-23* allele. The section shows *let-23* (ZK1067.1A.1) on chromosome II. Insertions that were made to generate *let-23(zh131)* are color coded; black: genomic sequence, red: FRT sites, orange: mutation of PAM for sgRNA3_let-23tyrK, blue: linker sequence, green: ATG of GFP coding region, yellow: Flag Tag.

